# Horizontal gene transfer by natural transformation promotes both genetic and epigenetic inheritance of traits

**DOI:** 10.1101/596379

**Authors:** Ankur B. Dalia, Triana N. Dalia

**Affiliations:** Department of Biology, Indiana University, Bloomington, IN 47405.

## Abstract

Natural transformation (NT) is a major mechanism of horizontal gene transfer in microbial species that promotes the spread of antibiotic resistance determinants and virulence factors. Here, we develop a cell biological approach to characterize the spatial and temporal dynamics of homologous recombination during NT in *Vibrio cholerae*. Our results directly demonstrate (1) that transforming DNA efficiently integrates into the genome as single-stranded DNA, (2) that the resulting heteroduplexes are resolved by chromosome replication and segregation, and (3) that integrated DNA is rapidly expressed prior to cell division. We show that the combination of these properties results in the epigenetic transfer of gene products within transformed populations, which can support the transgenerational epigenetic inheritance of antibiotic resistance in both *V. cholerae* and *Streptococcus pneumoniae*. Thus, beyond the genetic acquisition of novel DNA sequences, NT can also promote the epigenetic inheritance of traits during this conserved mechanism of horizontal gene transfer.

## Introduction

Microbes can share genetic material with one another via a variety of mechanisms that are cumulatively termed horizontal gene transfer (HGT). One broadly conserved mechanism for HGT is natural transformation (NT), which is a process whereby some microbial species can take up free DNA from the environment and subsequently integrate it into their genome by homologous recombination. This process is clinically relevant because NT is a property of many bacterial pathogens and can promote the acquisition and spread of antibiotic resistance determinants and virulence factors (Blokesch, 2016; Lorenz and Wackernagel, 1994). Despite 90 years of research since the discovery of NT (Griffith, 1928), many aspects of DNA integration during this process remain unclear.

During NT, cells generally interact with double-stranded DNA (dsDNA) in the environment, but only a single strand of this DNA is translocated into the cytoplasm (Gabor and Hotchkiss, 1966; Lacks, 1962; Piechowska and Fox, 1971). Upon uptake into the cytoplasm, this single-stranded DNA (ssDNA) is rapidly decorated by proteins including DprA and the recombination protein RecA (Kidane and Graumann, 2005; Mortier-Barriere et al., 2007). Prior work provides molecular evidence to suggest that this ssDNA can be directly integrated into the genome (Dubnau and Davidoff-Abelson, 1971; Fox and Allen, 1964; Mejean and Claverys, 1984). Thus, the ssDNA-RecA complex likely undergoes a homology search with the genome and RecA subsequently promotes strand invasion to generate a three-stranded D-loop intermediate.

Following strand invasion, the branches of the D-loop are then likely extended via a process known as branch migration. Indeed, recent work has identified natural transformation specific branch migration factors (Marie et al., 2017; Nero et al., 2018). Following branch migration, it is hypothesized that this three-stranded complex is then processed by an unresolved mechanism to generate a stable heteroduplex between the integrated transforming DNA (tDNA) and the genome. This heteroduplex could then be subjected to repair (e.g. via the mismatch repair system) or it is hypothesized that chromosome replication is required to resolve and segregate this heteroduplex. Direct evidence for many of the steps in this model of recombination, however, remains lacking.

Here, we develop and employ novel cell biological markers to test this model and extend our mechanistic understanding of the spatial and temporal dynamics of DNA integration during NT in single cells. Furthermore, this analysis has revealed that beyond the actual horizontal transfer of DNA, NT promotes an unappreciated mechanism for epigenetic inheritance that has broad implications for the spread of traits by this mode of horizontal gene transfer.

## Results

### ComM fluorescent fusions serve as a marker for homologous recombination during NT

There are currently no tools available to determine when cells are actively undergoing homologous recombination during NT, which has limited our understanding of this process. Fluorescent fusions to RecA do not fulfill this need because RecA interacts with ssDNA immediately upon uptake (Kidane and Graumann, 2005; Seitz and Blokesch, 2014); thus, making it difficult to distinguish between uptake of ssDNA into the cytoplasm from active integration into the chromosome. Also, recent work highlights the use of fluorescently labeled DNA to track the progress of DNA uptake and integration during NT (Boonstra et al., 2018; Corbinais et al., 2016); however, this approach suffers from the same shortfall. Recent work from our group has demonstrated that ComM is a hexameric helicase required for branch migration during NT in *V. cholerae* (Nero et al., 2018). We found that purified ComM only formed hexamers *in vitro* in the presence of ATP and ssDNA (Nero et al., 2018). Thus, we hypothesized that ComM may form active multimeric complexes specifically at the site of tDNA integration *in vivo*. To test this, we assessed the localization of a functional fluorescent fusion of ComM (Nero et al., 2018) in a constitutively competent strain of *V. cholerae* that exhibits high rates of natural transformation (transformation frequencies of up to ~50%) (Ellison et al., 2018). In the absence of any tDNA, GFP-ComM remained diffusely localized in the cytoplasm. In the presence of tDNA, however, GFP-ComM formed transient foci (Fig. 1A-C). We hypothesized that these foci represented the site of homologous recombination; specifically, sites of branch migration downstream of RecA-mediated strand invasion. Consistent with this, we did not observe any GFP-ComM foci in Δ*recA* or Δ*dprA* mutant backgrounds when cells were incubated with tDNA.

**Fig. 1.**
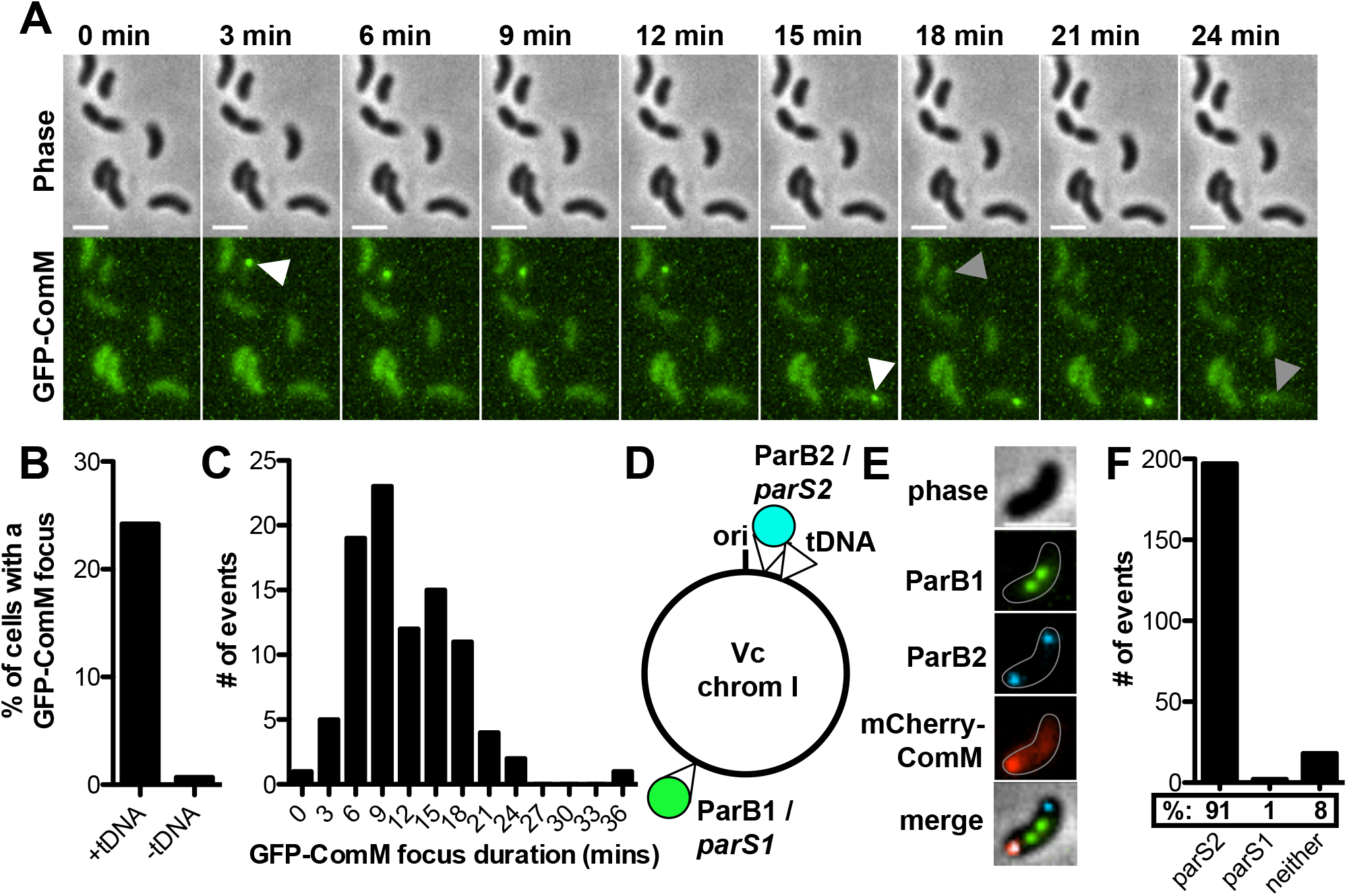
ComM fluorescent fusions serve as a marker for homologous recombination during NT. (**A**) Montage of timelapse imaging of a constitutively competent strain expressing GFP-ComM incubated with tDNA. White arrows indicate the appearance of GFP-ComM foci, while gray arrows indicate their disappearance. Scale bar, 2µm. (**B**) The percentage of cells that form a ComM focus when cells were imaged in the presence or absence of tDNA. Experiments were imaged every 1 min for 45 mins. Data are from three independent experiments and the mean is reported; n = 1315 cells for +tDNA and n = 1143 for -tDNA samples. (**C**) The duration of GFP-ComM foci from cells incubated with tDNA. n = 110 foci analyzed. (**D**) Schematic for the experimental setup to test whether ComM foci mark the site of homologous recombination. Cells constitutively expressed yGFP-ParB1/CFP-ParB2, contained a *parS2* site proximal to the origin, a *parS1* site proximal to the terminus, and expressed mCherry-ComM. Cells were incubated with tDNA that should integrate proximal to the *parS2* site. (**E**) Static image of a predivisional cell showing that the mCherry-ComM focus colocalizes with the proximally located *parS2* site. Scale bar, 2µm. (**F**) Quantifying colocalization of mCherry-ComM foci with the indicated *parS* site (i.e. ParB focus) when incubated with tDNA that integrates proximal to the *parS2* site. Data are from two independent experiments; n = 217 ComM foci analyzed.

Next, we wanted to further test whether ComM foci represented the site of homologous recombination during NT. To that end, we used two orthologous ParB/*parS* systems (Nielsen et al., 2006) to mark distinct locations of the *V. cholerae* genome. Cells in this assay constitutively expressed two distinct fluorescent ParB fusion proteins (yGFP-ParB1 and CFP-ParB2), which can specifically bind to their corresponding *parS* sites (*parS1* and *parS2*, respectively). Binding of ParB to *parS* forms a fluorescent focus that demarcates the location of the chromosomal *parS* locus within the cell. A *parS1* site was integrated proximal to the terminus, while a *parS2* site was integrated proximal to the origin. *V. cholerae* adopts a longitudinal ori-ter configuration during cell division (David et al., 2014; Fogel and Waldor, 2006), thus, this labeling scheme provides maximal spatial separation between the two *parS* foci. Next, we incubated cells with tDNA that would integrate in close proximity to (~10kb away from) the *parS2* site (Fig. 1D). If ComM foci formed at the site of homologous recombination, we would expect fluorescent ComM foci to colocalize with the *parS2* focus. Indeed, ~91% of the time, ComM foci colocalized with the *parS2* site (Fig. 1E-F). The few events where ComM foci formed distal to the *parS2* site (Fig. S1A) may represent attempts at illegitimate recombination, formation of ComM foci on tDNA independent of recombination, and/or a local loss of DNA compaction which results in spatial separation of the ComM and *parS2* foci. In some instances, ComM foci exhibited colocalized movement with the *parS2* site (Fig. S1B), which further supported the hypothesis that ComM foci formed at the site of DNA integration. Also, we performed the reciprocal experiment by incubating cells with tDNA that would integrate in close proximity (~10kb away from) to the *parS1* site. As expected, ComM foci colocalized with the *parS1* site ~90% of the time (Fig. S1C-E). Cumulatively, these results indicate that fluorescent ComM foci demarcate the site of homologous recombination during NT. Furthermore, these data establish that cell biological tracking of ComM and orthologous ParB/*parS* systems provides a robust spatial and temporal readout for homologous recombination in single cells.

### Tracking ComM allows for quantification of the success rate for homologous recombination during NT

It is currently unclear how efficacious the process of homologous recombination is during NT (i.e. how often does homologous recombination succeed after the process is initiated?). In fact, this is relatively poorly understood in most models of recombination. Above, we demonstrated that the formation of ComM foci at the site of DNA integration likely represent independent attempts at tDNA integration downstream of RecA-mediated strand invasion. These ComM foci do not, however, indicate whether these attempts were ultimately successful at integrating DNA or not. We hypothesized that if we could quantify the cells where integration succeeded, we would have a powerful metric to assess the efficacy of integration during NT. To assess successful tDNA integration, we used a strain that constitutively expressed CyPET-ParB1 and initially lacked a cognate *parS1* site, which resulted in diffuse CyPET-ParB1 localization. We then transformed these cells with tDNA that would integrate a *parS1* site into the genome. Upon successful tDNA integration, a *parS1* site would be formed resulting in the formation of a CyPET-ParB1 focus. These cells also expressed a GFP-ComM fusion. Thus in this experiment, we could assess attempts at tDNA integration by tracking the formation of GFP-ComM foci and identify cells where tDNA integration succeeded by tracking the formation of CyPET-ParB1 foci.

First, we incubated cells with tDNA that would insert a 148bp *parS1* site into the genome (i.e. Δ0kb::*parS1*). We found that ~80% of cells generated at least one ComM focus during the duration of the experiment (Fig. 2A-C); however, only a subset of these cells produced a ParB1 focus (Fig. 2A), indicating that many attempts (as assessed by ComM foci) ultimately failed to successfully integrate tDNA. Based on this, we could calculate a success rate for tDNA integration. The success rate on a per cell basis, defined as the percentage of cells that attempted to integrate tDNA (i.e. formed at least 1 ComM focus) that ultimately succeeded (i.e. formed a ParB1 focus) was ~45% (Fig. 2D). However, many cells produced more than one ComM focus throughout the duration of the experiment (Fig. 2A-C). If we assume that only a single ComM focus is required for successful tDNA integration we can also calculate the success rate on a per attempt (i.e. per focus) basis. This is defined as the percentage of cells that formed a ParB1 focus relative to the total number of ComM foci observed. The success rate per attempt observed was ~25% (Fig. 2D). Interestingly, there was no difference in the duration of GFP-ComM foci when we compare cells that ultimately succeeded vs failed to integrate tDNA (Fig. 2E). This suggests that the duration of the ComM-dependent branch migration complex does not predict successful integration. We did, however, observe a slight increase in the number of ComM foci produced in cells that ultimately succeeded to integrate tDNA (Fig. 2F).

**Fig. 2.**
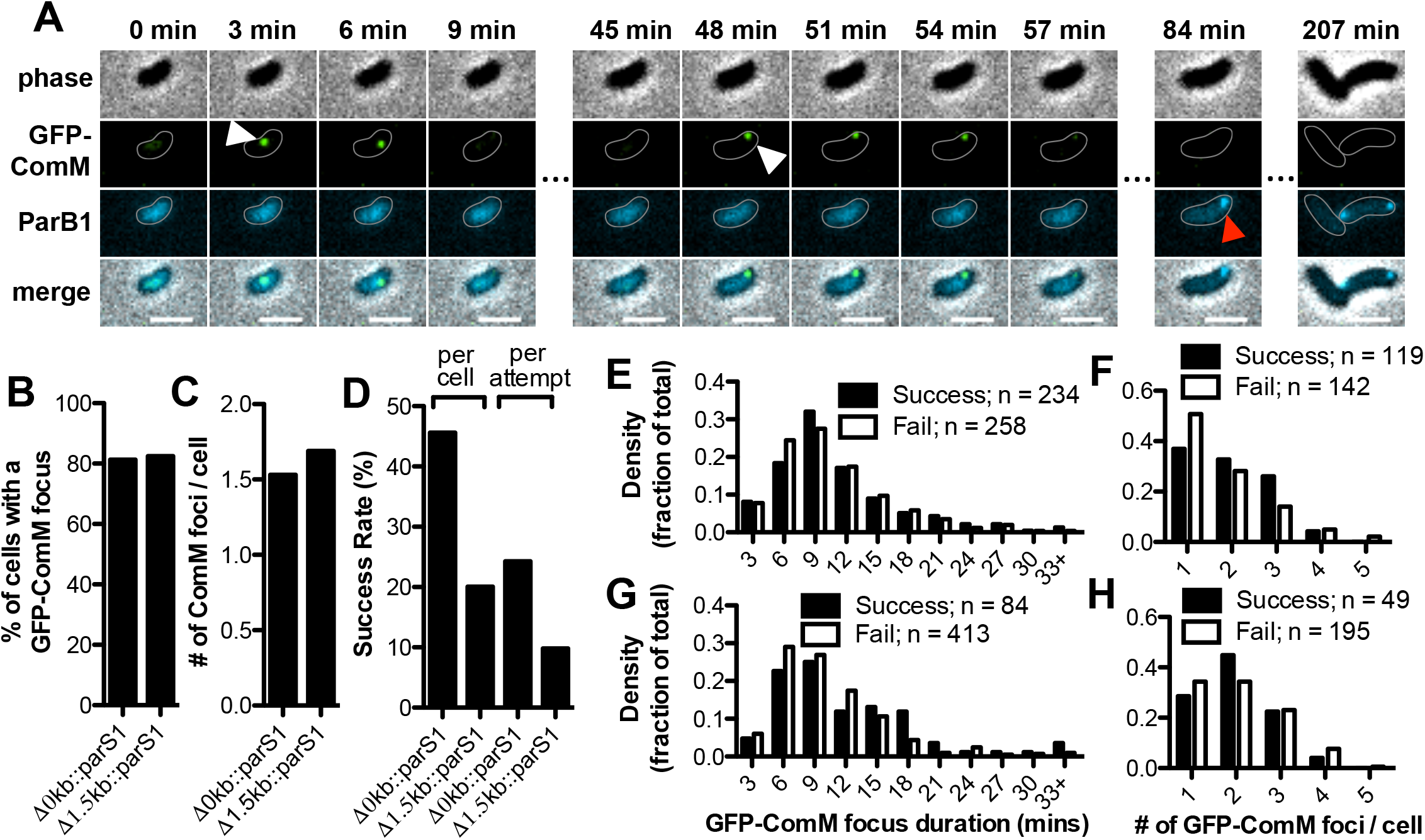
Quantifying the efficacy of homologous recombination during NT. Success rates for homologous recombination were determined using a cell biological approach where cells expressed GFP-ComM and CyPet-ParB1, but initially lacked a *parS1* site. These cells were then incubated with tDNA that would simply integrate a *parS1* site into the genome (i.e. Δ0kb::*parS1*) or delete and replace a 1.5 kb marker with a *parS1* site (i.e. Δ1.5kb::*parS1*). (**A**) Montage of timelapse imaging of one representative example of successful tDNA integration for Δ0kb::*parS1*. White arrows show when the cell forms ComM foci indicating an attempt at tDNA integration, while red arrows show when the cell forms a CyPet-ParB1 focus, which indicates successful tDNA integration into the genome. Experiments were imaged every 3 min for 5 hours. (**B**) The percentage of cells that formed at least one ComM focus and (**C**) the number of ComM foci per cell. n = 320 for Δ0kb::*parS1* and n = 296 for Δ1.5kb::*parS1*. (**D**) The success rate for tDNA integration was calculated on a per cell basis (i.e. the percentage of cells that formed ComM foci, which ultimately succeeded to integrate tDNA) or a per attempt basis (i.e. assuming that only one ComM focus is required for integration, the success rate per attempt is the percentage of ComM foci that resulted in successful tDNA integration). n = 261 cells with ≥1 ComM focus analyzed for Δ0kb::*parS1* and n = 244 for Δ1.5kb::*parS1*. Histograms showing ComM focus duration in cells that ultimately succeeded vs failed to integrate tDNA for (**E**) 0kb::*parS1* and (**G**) Δ1.5kb::*parS1*. Histograms showing the number of ComM foci in cells that succeeded vs failed to integrate tDNA for (**F**) 0kb::*parS1* and (**H**) Δ1.5kb::*parS1*.

The transformation frequency for introducing large insertions and deletions by natural transformation is lower than when integrating smaller mutational cargo (Dalia et al., 2014). If ComM foci truly represent attempts at DNA integration, we hypothesized that when using tDNA that would introduce a large deletion, that the number of ComM foci formed (i.e. the number of attempts) would remain unchanged, while the success rate would be reduced. To test this, we performed an assay using cells where the targeted integration site already contained a ~1.5kb chloramphenicol resistance (Cm^R^) cassette. Thus, integration of the *parS1* site would require cells to delete the 1.5kb Cm^R^ cassette and replace it with the *parS1* site (i.e. Δ1.5kb::*parS1*), while the cells used in the prior experiment merely had to insert the parS1 site into the genome (i.e. Δ0kb::*parS1*). For Δ1.5kb::*parS1*, we found that cells generated just as many ComM foci as for Δ0kb::parS1, indicating that cells were attempting to integrate tDNA at the same rate (Fig. 2A-C). Importantly, the arms of homology in the tDNA were identical in both experiments. Thus, this result is consistent with RecA-mediated strand invasion in one arm of homology occurring at equal efficiencies regardless of the nature of the mutation being introduced. The duration of ComM foci and the number of foci generated per cell were not markedly different between Δ1.5kb::*parS1* and Δ0kb::*parS1* (Fig. 2E-H). The success rate, however, was approximately half for Δ1.5kb::*parS1* compared to Δ0kb::*parS1* (Fig. 2D). Thus, deletion of ~1.5kb results in an ~2-fold reduction in the success rate for integration. Because the number of attempts at DNA integration were equivalent between Δ1.5kb::*parS1* and Δ0kb::*parS1*, the success rate calculated here is indicative of the processes that must occur downstream of RecA-mediated strand invasion, which include ComM-dependent branch migration.

In both of the experiments described above, formation of the ParB1 focus (i.e. the indicator for successful tDNA integration) was temporally delayed from the formation of ComM foci.

Integration of ssDNA, as hypothesized for NT, might account for this delay because a single-stranded *parS* site is not sufficient to bind to parB (Pillet et al., 2011). Thus, we would expect that a ParB1 focus would only form after the *parS1* site is converted into dsDNA (e.g. via chromosome replication). Consistent with this hypothesis, we consistently observed that only one of the two daughter cells inherited a ParB1 focus, and in the subsequent round of division both daughter cells inherited a ParB1 focus (Fig. 2A). We therefore hypothesized that tDNA integrates as ssDNA and the resulting heteroduplex is resolved by chromosome replication, which we test further below.

### tDNA integrates as ssDNA during NT to form a heteroduplex that is resolved by chromosome replication

Decades of research has provided molecular evidence that tDNA likely integrates into the genome as ssDNA during NT (Dubnau and Davidoff-Abelson, 1971; Fox and Allen, 1964; Mejean and Claverys, 1984). To provide more direct and quantitative evidence for this, we started with cells that constitutively expressed yGFP-ParB1/CFP-ParB2 and contained a single *parS2* site. We then transformed these cells with tDNA that would replace the *parS2* site with a *parS1* site. If ssDNA integrates during NT, then we would expect chromosome replication to generate two genomes, where one has a *parS1* site and the other has a *parS2* site. This would be observed experimentally as predivisional cells that contain both green ParB1 and cyan ParB2 foci (Fig. 3A). Alternatively, if tDNA is integrated as dsDNA into the genome, we should only observe cells that exclusively have ParB1 or ParB2 foci (Fig. S2A). When we performed this experiment, we found that ~94% of cells exhibited a phenotype consistent with the integration of ssDNA (Fig. 3B-C, Fig. S2B, and **movie S1**). For cells that exhibited ssDNA integration, we found that cells faithfully replicated their chromosomes in the next round of division (i.e. the daughter cell that inherited the *parS1* site in the first round of division then replicated to form two *parS1* foci in the subsequent round of chromosome replication) (Fig. 3B). The observation of dsDNA integration in ~6% of cells may be the result of uptake and integration of multiple tDNA molecules that replace both the leading and lagging strands of the endogenous genome. This is not unprecedented because it is well established that naturally transformable species can take up and integrate multiple independent tDNA molecules in a process termed congression or cotransformation (Dalia et al., 2014; Nester et al., 1963). These data indicate that during NT, cells predominantly integrate ssDNA into the genome and that the resulting heteroduplex is resolved by chromosome replication and segregation.

**Fig. 3.**
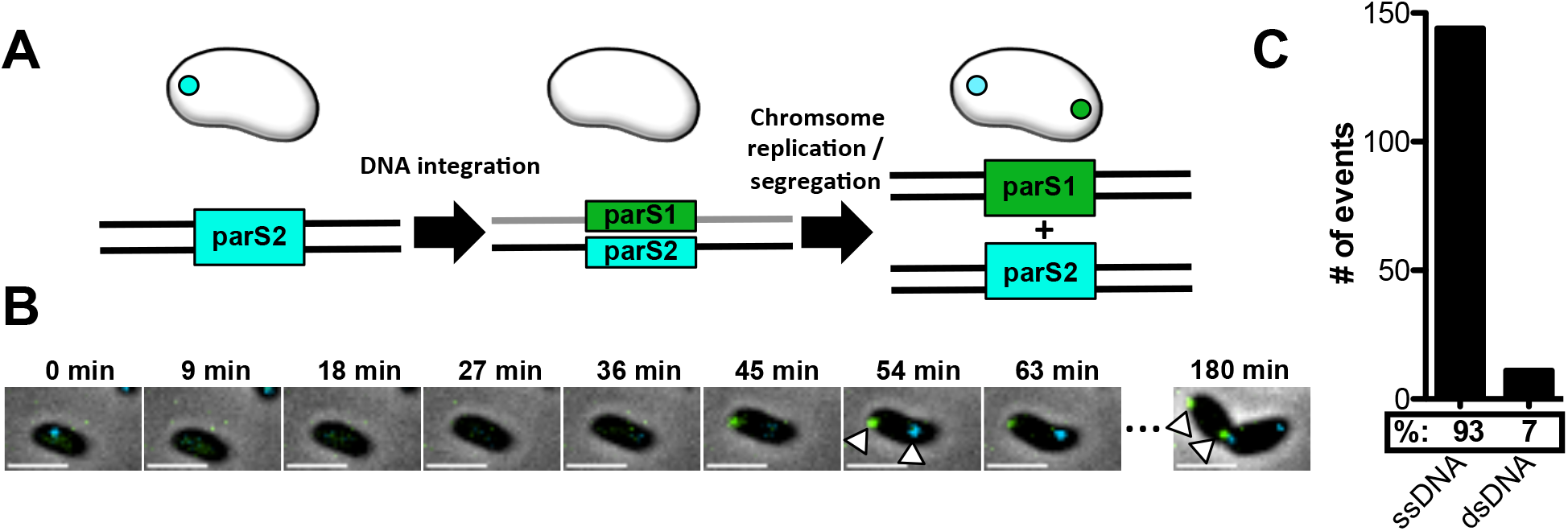
Integration of single-stranded tDNA during NT forms heteroduplexes that are resolved by chromosome replication and segregation. (**A**) Schematic of the experimental setup and expected results if tDNA is integrated as ssDNA during NT. Cells constitutively expressed yGFP-ParB1/CFP-ParB2 and contained a *parS2* site (cyan ParB2) in the genome. Cells were then transformed with tDNA that would replace the *parS2* site with a *parS1* site (green ParB1). Experiments were imaged every 3 min for 5 hours. (**B**) Montage of timelapse imaging for one representative example of ssDNA integration. Arrows indicate a predivisional cell with both yGFP-ParB1 and CFP-ParB2 foci, which indicates that the DNA that was integrated during NT was single-stranded and the resulting heteroduplex was resolved by chromosome replication and segregation. Consistent with these new foci representing a resolved heteroduplex, each *parS* site is faithfully replicated in the next round of chromosome replication (arrows at 180 min). Scale bars, 2µm. (**C**) Quantification of the number of cells that exhibit ssDNA integration (as indicated in **B**) vs dsDNA integration (as indicated in Fig. S3). Data are from three independent experiments; n = 155 cells with integrated tDNA analyzed.

### ComM-dependent branch migration immediately precedes DNA integration during NT

Previously, we used insertion of a *parS1* site to identify cells that successfully integrated DNA (Fig. 2). Because cells integrate ssDNA, however, they only generated a ParB1 focus after chromosome replication, which delayed the readout for successful integration from the time that integration actually occurred. To test whether we could generate a more sensitive temporal indicator of tDNA integration, we used a modified experimental approach by assessing the disruption of an established *parS* site (Fig. S3A). We started with cells that constitutively expressed yGFP-ParB1/CFP-ParB2 and harbored *parS1* and *parS2* sites in close proximity (~10kb apart) on the chromosome, with the *parS1* site disrupting a *gfp* gene. These strains also contained an mCherry-ComM fusion expressed at the native locus. We then transformed these cells with tDNA that would delete the *parS1* site and restore the *gfp* gene. Thus, we can assess the spatial and temporal relationship between ComM activity (i.e. mCherry-ComM foci) and tDNA integration (i.e. loss of the ParB1 focus). The presence of a constant ParB2 focus served as a control for local rearrangements of the genome (which should disrupt both ParB1 and ParB2 foci), while specific tDNA integration should only result in loss of the ParB1 focus. As a secondary indicator of successful tDNA integration (beyond the loss of the ParB1 focus), transformed cells should express high levels of GFP above the baseline fluorescence of the yGFP-ParB1 construct once the *gfp* gene is restored. When we performed this experiment, we found that loss of the ParB1 focus was immediately preceded by the formation of a colocalized ComM focus (Fig. S3B). ParB1 then remained diffusely localized until chromosome replication, as evidenced by the splitting of the ParB2 focus, whereupon we saw reappearance of one ParB1 focus (Fig. S3B). This is consistent with ssDNA integration of tDNA into the genome because chromosome replication would be expected to restore the *parS1* site from the DNA strand that was not replaced (Fig. S3A). Following chromosome replication, GFP expression increased, which further confirmed that cells were transformed (Fig. S3B). These results indicate that DNA integration occurs rapidly following the formation of ComM foci and that monitoring the loss of an established *parS* site (i.e. a ParB focus) provides the most immediate readout for homologous recombination during NT.

### Expression of integrated tDNA occurs rapidly after chromosome replication and prior to cell division

The experiments described above established the spatial and temporal link between ComM dependent branch migration and tDNA integration. Assessing the exact timing of integrated tDNA in these experiments, however, was complicated by the fact that cells already expressed a baseline amount of yGFP-ParB1 (Fig S3B). This made it difficult to determine the exact timing of expression for the repaired *gfp* gene following DNA integration because it was assessed in the same fluorescent channel as yGFP-ParB1.

To better define when integrated tDNA was expressed, we altered the experimental setup so that the expression of integrated tDNA was assessed in a distinct fluorescent channel (Fig. 4A). We generated strains that expressed mCherry-ParB1/CFP-ParB2 and harbored *parS1* and *parS2* sites in close proximity on the genome where the *parS1* site disrupted a *gfp* gene. Cells were then transformed with tDNA to delete the *parS1* site and restore the *gfp* gene (Fig. 4A). Due to a lack of available fluorescent channels, ComM was not tracked in these experiments; however, tracking ComM activity was not critical to assess the timing of tDNA expression. When we performed these experiments, we observed tDNA integration as the loss of the ParB1 focus and subsequent restoration of this focus upon chromosome replication (Fig. 4B, **movie S2**), which is consistent with ssDNA integration as observed previously (Fig. 3B). This analysis also revealed that integrated tDNA was rapidly expressed following chromosome replication (Fig. 4B, **movie S2**). The delay to *gfp* expression following chromosome replication was ~15 mins (Fig. 4C), while the doubling time in these assays was ~3 hours. The maturation time for the *gfp* allele used is ~6.5 mins (Megerle et al., 2008). Thus, these results indicate that integrated tDNA is expressed almost immediately following chromosome replication, which resolves the integrated heteroduplex.

**Fig. 4.**
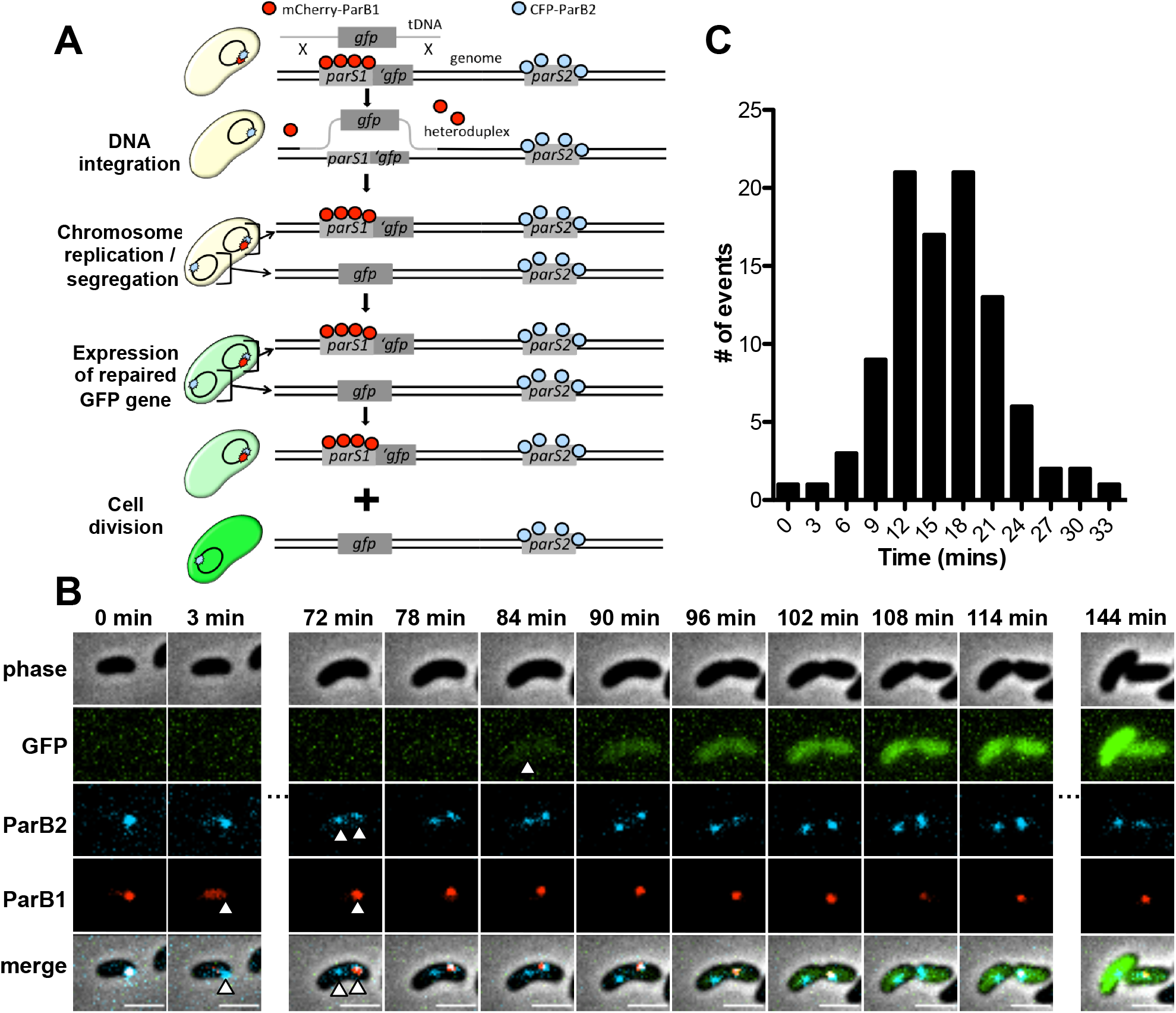
Natural transformation promotes epigenetic inheritance of tDNA-derived gene products. (**A**) Schema of the experimental setup. Cells contained *parS1* (red ParB1) and *parS2* (cyan ParB2) sites in close proximity on the chromosome. The *parS1* site interrupts a chromosomally integrated *gfp* gene. Cells were then transformed with tDNA to delete the *parS1* site and restore the *gfp* gene. Experiments were imaged every 3 mins for 5 hours. (**B**) Montage of time lapse imaging of one representative cell that successfully integrates tDNA. Cells first integrate DNA (as indicated by loss of the ParB1 focus at 3 min), then chromosome replication and segregation occurs to resolve the heteroduplex (as indicated by splitting of the ParB2 focus and reappearance of the ParB1 focus in one genome at 72 min), and finally GFP is rapidly expressed following chromosome replication (84 min timepoint) and inherited by both daughter cells, including the cell that is genotypically still a GFP mutant (i.e. the cell containing the ParB1 focus). Scale bar, 2 µm. (**C**) Histogram of the time delay to tDNA-derived GFP expression following chromosome replication. Data are from three independent experiments and n = 97 GFP positive cells analyzed.

### NT promotes transgenerational epigenetic inheritance of antibiotic resistance during NT

Because tDNA is integrated as ssDNA, the predivisional transformed cell has two genomes of distinct genotypes following chromosome replication – one where the *gfp* gene is still mutated (Fig. 4B – the genome represented by the colocalized ParB1/ParB2 foci) and one where the *gfp* gene is repaired (i.e. Fig. 4B - the genome represented by the independent parB2 focus). Expression of the integrated tDNA (from the restored *gfp* gene) in this predivisional cell occurred prior to cell division. Thus, despite the fact that one of the daughter cells is genotypically a *gfp* mutant, it still epigenetically inherits a significant amount of the GFP gene product derived from the tDNA integrated into its sibling.

While epigenetic inheritance of GFP may be inconsequential, there may be important contexts in which the epigenetic inheritance of tDNA-derived products can provide a benefit. One instance could be in the context of antibiotic resistance where epigenetic inheritance of an antibiotic resistance gene product could transiently protect the sibling of a transformed cell. We setup an experimental approach to directly test this hypothesis (Fig. S4A-B). We started with cells that had a chloramphenicol resistance marker (Cm^R^) integrated into their genome. We then transformed these cells with tDNA that would replace the Cm^R^ marker with a kanamycin resistance marker (Kan^R^). So, the genotype of cells in this experiment should theoretically be exclusively Cm^R^ or Kan^R^ (Fig. S4A). Upon tDNA integration and subsequent chromosome replication, we would expect predivisional cells to have 2 distinct genomes – one that is Cm^R^ and the other that is Kan^R^ (Fig. S4A). Because integrated tDNA is expressed rapidly following chromosome replication and prior to cell division, we expect that the Kan^R^ gene product should be inherited by both the Kan^R^ daughter cell and the Cm^R^ daughter cell. Thus, the genetically Cm^R^ siblings of transformed cells may be phenotypically resistant to kanamycin (Fig. S4A). To assess this, we challenged these transformation reactions with a lethal dose of kanamycin to kill susceptible cells (Fig. S4B). Cells were then washed to remove the kanamycin and plated for quantitative culture on selective plates to determine the number of genetically Kan^R^ and Cm^R^ cells that survived the kanamycin treatment (Fig. S4B). We also assessed the number of illegitimate recombinants (i.e. transformants where the Kan^R^ marker was integrated at a locus other than the intended site), which would be genetically Kan^R^+Cm^R^.

As expected, when cells were not incubated with any tDNA, very few (~10^3^) genetically Cm^R^ or Kan^R^ cells survived the kanamycin treatment (Fig. 5A-C, **0 min white bars**). When cells were incubated with Kan^R^ tDNA, there were ~10^7^ genetically Kan^R^ cells that survived the kanamycin treatment (Fig 5B, **0 min black bars**), which represents the number of genetic transformants generated (i.e. the number of cells that actually integrated the Kan^R^ tDNA into their genome). We also observed that a similar number of genetically Cm^R^ cells survived the kanamycin treatment (Fig. 5A, **0 min black bars**), and importantly, this was ~4 logs greater than the number of illegitimate recombinants observed (Fig. 5C, **0 min black bars**). The genetically Cm^R^ cells that survived the kanamycin treatment represent cells that are phenotypically Kan^R^, likely through epigenetic inheritance of the Kan^R^ gene product from their transformed sibling. Interestingly, of all cells that survived the kanamycin treatment, ~25% of these cells were Cm^R^ (Fig. 5D, **0 min**). The theoretical maximum if every transformed cell protected its sibling would be 50%. Thus, these results indicate that ~1 in every 2 transformants conveyed epigenetic kanamycin resistance to its untransformed sibling.

**Fig. 5.**
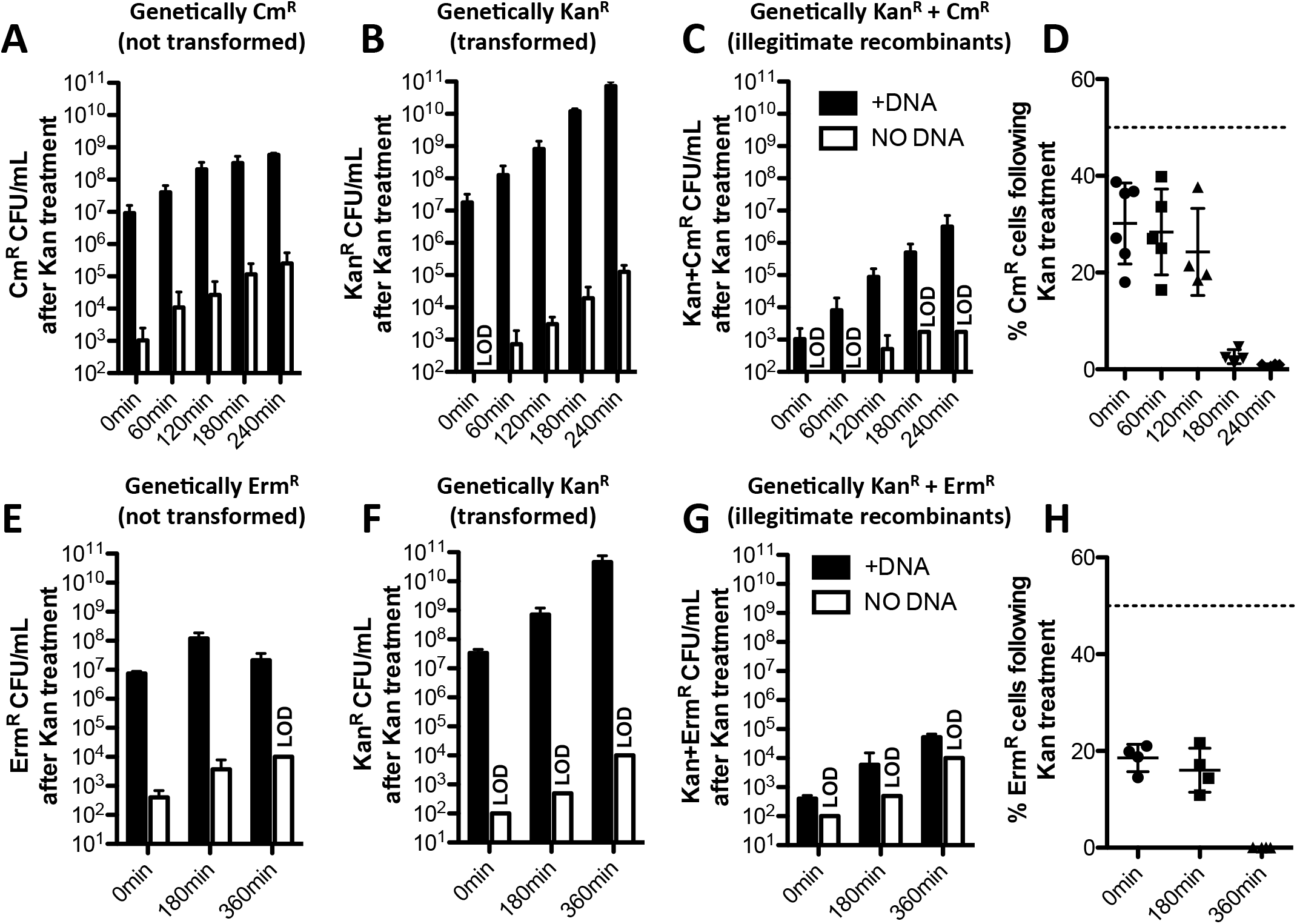
NT confers transgenerational epigenetic inheritance of antibiotic resistance in *V. cholerae* and *S. pneumoniae*. Transgenerational epigenetic inheritance of antibiotic resistance was assessed as schematized in Fig. S4A-B in (**A**-**D**) *V. cholerae* and (**E**-**H**) *S. pneumoniae*. In both cases, cells were transformed with Kan^R^ tDNA and then outgrown for the amount of time indicated prior to a brief treatment with a high dose of kanamycin to kill susceptible cells. Cells were then washed and plated for quantitative culture on the media indicated to quantify the number of (**A** and **E**) non-transformants, (**B** and **F**) transformants, and (**C** and **G**) illegtimate recombinants that survived the kanamycin treatment. (**D** and **H**) Of the cells that survived the kanamycin treatment, the percentage of cells that were non-transformants was calculated from **A**-**B** and **E**-**F** and plotted. The dotted line at 50% represents the theoretical maximum value if every transformant conferred phenotypic kanamycin resistance to its untransformed sibling. All data are shown as the mean ± SD and are from at least 4 independent biological replicates.

As noted above, the phenotypic kanamycin resistance observed in genetically Cm^R^ cells is likely the result of epigenetic inheritance of the Kan^R^ gene product. Because these cells do not have the capacity to continue to produce the Kan^R^ gene product, we would expect that the gene product and resulting phenotypic kanamycin resistance should dilute out in these cells through growth. So next, we wanted to determine how many generations phenotypic kanamycin resistance was maintained in genetically Cm^R^ cells. To test this, we performed the same experiment described above, however, we outgrew reactions in rich medium for 60, 120, 180, and 240 mins prior to subjecting them to the kanamycin treatment (Fig. S4B). Surprisingly, we found that phenotypic kanamycin resistance was maintained in some portion of Cm^R^ cells even out to 240 mins of outgrowth, which represents ~11 generations of growth (Fig. 5A). We did, however, see that the percentage of cells that were Cm^R^ following the kanamycin treatment dropped significantly after 180 mins of outgrowth (Fig. 5D). But this ratio was consistently maintained for 120 mins of outgrowth, which represents ~5-6 generations of growth. Thus, these data indicate that the epigenetic inheritance of phenotypic kanamycin resistance is maintained for an extended number of generations.

To determine if integration of tDNA was required to confer epigenetic kanamycin resistance we performed this assay in Δ*recA* and Δ*comM* mutant backgrounds. The Δ*recA* strain, as expected, did not produce any Kan^R^ transformants and there was no phenotypic kanamycin resistance observed in Cm^R^ cells (Fig. S4C). Prior work, however, indicates that tDNA is less stable in the cytoplasm of Δ*recA* mutants (Berge et al., 2003). The Δ*comM* strain on the other hand has an ~100-fold lower transformation frequency compared to the parent, while DNA uptake and stability of tDNA in the cytoplasm remains unchanged (Nero et al., 2018). In the Δ*comM* mutant, we observed an ~100-fold reduction in the number of transformants (i.e. Kan^R^ cells) and a corresponding decrease in the number of Cm^R^ cells that survived the kanamycin treatment (Fig. S4C), which is consistent with tDNA integration being required for epigenetic inheritance of kanamycin resistance. Transgenerational epigenetic inheritance of antibiotic resistance is not unique to Kan^R^ because similar results were obtained when using tDNA containing a spectinomycin resistance cassette (Spec^R^) and challenging cells with a lethal dose of spectinomycin (Fig. S4D).

We also tested epigenetic inheritance of antibiotic resistance in the model Gram-positive naturally transformable species *Streptococcus pneumoniae*. Transformation of *S. pneumoniae* with Kan^R^ tDNA conferred transgenerational phenotypic kanamycin resistance to untransformed cells (Fig. 5E-H), which likely occurred through epigenetic inheritance of the Kan^R^ gene product as observed in *V. cholerae* (Fig. 5A-D).

Other antibiotics/resistance determinants could not be tested using the experimental approach described above due to a requirement for the antibiotic to exhibit bactericidal activity following a high dose, short duration exposure. Therefore, we sought to establish a microscopy-based assay that would allow us to directly observe epigenetic inheritance of antibiotic resistance in single cells, which would be amenable to testing non-bactericidal antibiotics. To that end, we generated strains that constitutively expressed mCherry-ParB1/CFP-ParB2 and had a single *parS1* site disrupting a *gfp* gene (i.e. *gfp*::*parS1*). These cells were then transformed with tDNA that would simultaneously remove the *parS1* site, repair the *gfp* gene, and integrate a *parS2* site linked to an antibiotic resistance marker (Ab^R^) (Fig. S5A). Thus, following tDNA integration and subsequent chromosome replication, predivisional transformed cells would have one genome that represents the parent genotype (i.e. the untransformed sibling = *gfp*::*parS1*, Ab^S^), while the other genome will represent the transformed genotype (i.e. the transformed sibling = *parS2*, *gfp* intact, and Ab^R^). Based on our previous results, we would expect the untransformed sibling to epigenetically inherit tDNA-derived gene products from its transformed sibling, which would allow it to grow on the antibiotic despite the fact that it does not actually encode an Ab^R^ marker. Indeed, this was exactly what we observed using tDNA containing a Kan^R^ marker (Fig. 6, **movie S3**), which also validated the results described above (Fig. 5A-D). Furthermore, when we performed this experiment using tDNAs that conferred resistance to other antibiotics (erythromycin and chloramphenicol), similar results were obtained (Fig. S5B-C, **movies S4-S5**), indicating that transgenerational epigenetic inheritance of antibiotic resistance during NT is not limited to bactericidal aminoglycosides like kanamycin and spectinomycin.

**Fig. 6.**
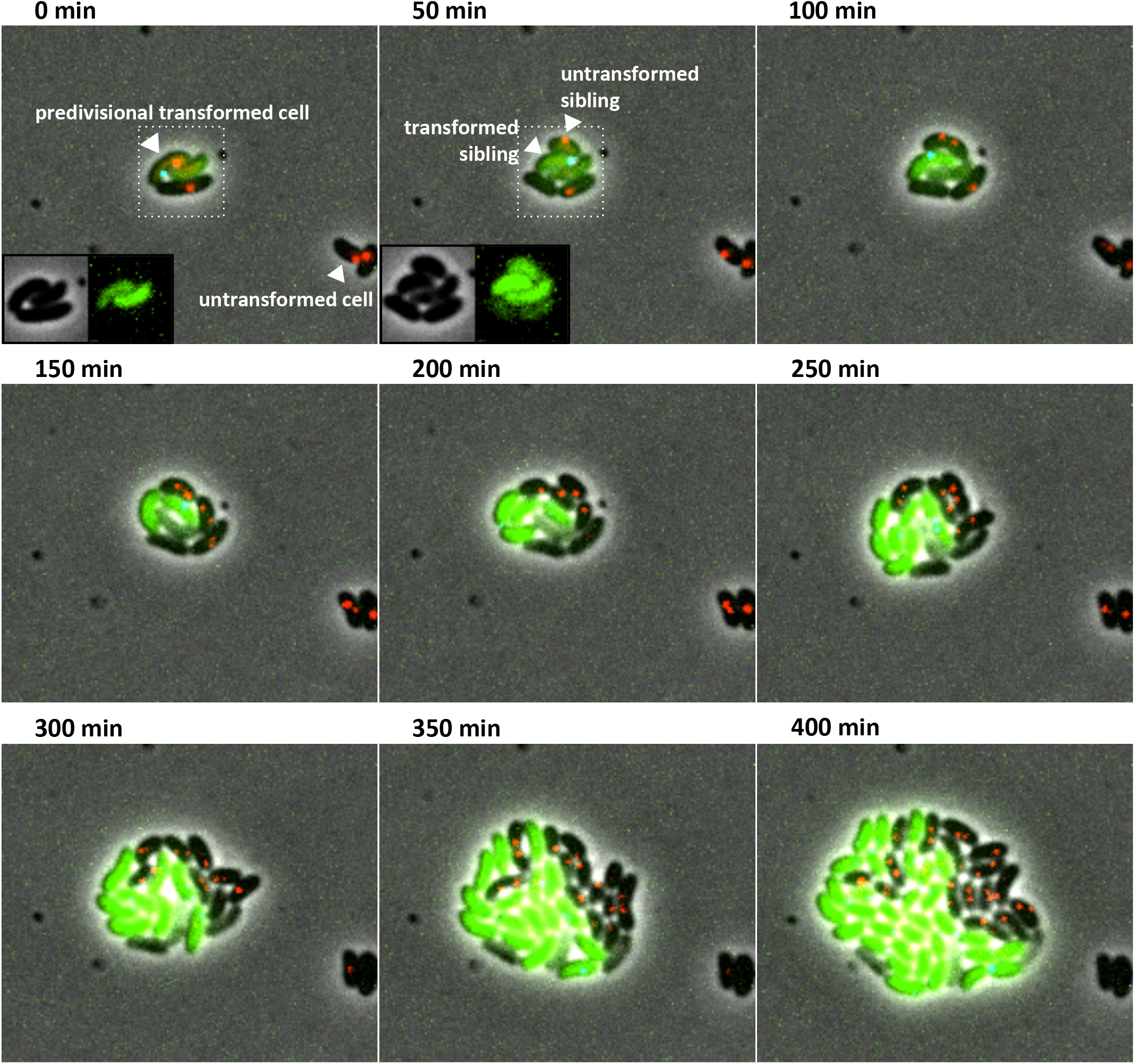
Single cell assay to demonstrate transgenerational epigenetic inheritance of kanamycin resistance during NT. For a detailed schema of the experimental setup see Fig. S5A. The parent strain constitutively expressed mCherry-ParB1 (red)/CFP-ParB2 (cyan) and contained a *parS1* site in the genome that disrupts a *gfp* gene. Cells were then incubated with tDNA that would remove the *parS1* site, restore the *gfp* gene, and integrate a *parS2* site linked to a Kan^R^ marker. Thus, transformed cells (i.e. cells that actually integrated the tDNA), would express GFP (green), contain a CFP-ParB2 focus, lack an mCherry-ParB1 focus, and should express the Kan^R^ gene product. While untransformed cells should represent the parent genotype and should not express GFP, contain an mCherry-ParB1 focus, lack a CFP-ParB2 focus, and should not express Kan^R^. Cells representing the parent genotype that were siblings of a transformed cell (i.e. the “untransformed sibling”), however, could epigenetically inherit the GFP and Kan^R^ gene products. Montage of timelapse imaging of one representative predivisional transformed cell under an M9+glucose pad containing 100 µg/mL kanamycin is shown. Early timepoints indicate that the untransformed sibling inherits GFP that was expressed prior to cell division, which serves as a proxy for the Kan^R^ gene product (see insets at 0 min and 50 min for images of just the GFP channel). As expected, the transformed sibling, which is genetically Kan^R^, grows well in the presence of kanamycin. Compared to the completely untransformed cell, which does not grow, the untransformed sibling (which is genetically Kan^S^) grows and divides for a number of generations in the presence of kanamycin, which indicates transgenerational epigenetic inheritance of the Kan^R^ gene product. Experiments were imaged every 5 min for 10 hours.

## Discussion

Our results provide direct evidence to validate the current model of tDNA integration during NT. Furthermore, they have provided invaluable insight into the spatial and temporal dynamics during the integration process at single cell resolution, which ultimately revealed an unappreciated mechanism for epigenetic inheritance during NT. Specifically, we provide direct and quantitative evidence that tDNA integrates as ssDNA into the genome. These results support decades of prior molecular work (Dubnau and Davidoff-Abelson, 1971; Fox and Allen, 1964; Mejean and Claverys, 1984). We demonstrate that following integration of ssDNA that the resulting heteroduplex is resolved by chromosome replication and segregation, which forms predivisional cells harboring genomes of two distinct genotypes (one containing the integrated DNA and the other lacking it). We also found that integrated tDNA is rapidly expressed immediately following chromosome replication, which occurs before cell division under the conditions tested. Importantly, we show that the combination of these properties - ssDNA integration, resolution of heteroduplexes by chromosome replication, and the rapid expression of tDNA in predivisional cells – allows for the epigenetic transfer of tDNA derived gene products to the untransformed siblings of transformed cells. Here, we observed evidence for epigenetic inheritance during NT in both *V. cholerae* and *S. pneumoniae*. Thus, it is likely that the mechanism for epigenetic inheritance described here is broadly applicable to this mode of horizontal gene transfer. In support of this, recent studies have shown that the expression of integrated DNA prior to cell division likely also occurs in naturally transformable *Helicobacter pylori* and *Bacillus subtilis* (Boonstra et al., 2018; Corbinais et al., 2016).

Our results demonstrate that fluorescent fusions of the branch migration factor ComM serve as a novel cell biological marker for homologous recombination during NT. Using this tool, we were able to quantify the success rate for DNA integration during NT *in vivo* as ~20-45% depending on the nature of the mutation being introduced. HGT by Hfr conjugation in *E. coli* was previously estimated to occur at an efficiency of ~96% (Babic et al., 2008). This was measured microscopically using conjugative transfer between Dam+ Hfr donors and Dam-recipients containing SeqA-YFP, which binds and forms fluorescent foci on the hemi-methylated DNA that forms in recipients. If SeqA-YFP foci persisted for >4 hours, it was assumed that DNA was stably integrated into the genome. Our analysis on the success rates for homologous recombination during NT required integration of a defined DNA cargo (a *parS* site), while the prior analysis of HGT by Hfr was sequence-independent (Babic et al., 2008)(recombination of any region of the methylated Hfr donor DNA should still yield a seqA-YFP focus in the recipient). Thus, it is entirely possible that integration of donor DNA within the homology arms of tDNA in our assays occurred at a higher frequency than what was observed for integration of the *parS* site. However, we cannot currently estimate this using the same approach outlined above for *E. coli* because Dam is essential for viability in *V. cholerae*. The readouts we developed for attempts (ComM foci) and successful homologous recombination (integration of a *parS* site) during NT, however, provide a sensitive assay to measure the effects of different mutational cargo (insertions vs deletions) on the rates of homologous recombination. Because we observed similar rates of attempts for integration of different tDNAs (i.e. the number of ComM foci) despite differing rates of successful integration, this analysis also demonstrates that RecA-dependent homology searching / strand invasion are not affected by the mutational cargo because these steps precede the formation of ComM foci.

It remains formally possible that epigenetic inheritance of gene products during NT is a mere byproduct of the integration process and does not confer any meaningful evolutionary advantage. We have, however, provided experimental evidence and further discussion below to highlight some potential advantages to this mode of epigenetic inheritance during NT. Experimentally, we demonstrate that epigenetic inheritance during NT can have important biological consequences for transformed populations using antibiotic resistance as an example. Specifically, we show that upon integration of tDNA encoding for an antibiotic resistance determinant that both the transformed cell and its untransformed sibling are both phenotypically resistant to the antibiotic. Perhaps most surprisingly, we found that this epigenetic inheritance of antibiotic resistance was long lasting and could confer resistance to untransformed cells for multiple generations. While we only observed growth of the untransformed cells for 3-5 generations in the presence of the antibiotics (Fig. 6, Fig. S5B-C, **movie S3-S5**), our antibiotic challenge assays suggest that tolerance to the drug may last significantly longer (Fig. 5). Natural environments are generally dynamic and exposure to stressors like antibiotics may only be transient. Thus, epigenetic inheritance during NT may provide an advantage in natural settings by providing transient resistance to these stressors. Interestingly, the resistance determinants used in this study act by different mechanisms (Kan^R^/Spec^R^ = inactivate drug, Cm^R^ = efflux pump, Erm^R^ = modifies the drug target), however, all are enzymatic mechanisms of resistance. If cells inherit even small amounts of these enzymes they may confer phenotypic drug resistance, which could account for why resistance is maintained for multiple generations. The transgenerational epigenetic inheritance of any traits via this mode of horizontal transfer, however, is subject to the expression, stability, and activity of the gene products that confer them.

There are three major modes of horizontal gene transfer that can modify bacterial genomes through recombination. These are natural transformation, conjugation, and phage transduction. Integration of ssDNA, which is required for the epigenetic transfer of traits by NT, is somewhat unique among these. Conjugation horizontally transfers a single strand of DNA into recipients; however, it is believed that the complementary strand is rapidly synthesized upon entering recipients prior to recombination (Willetts and Wilkins, 1984). Likewise, phage transduction can result from the injection of dsDNA or ssDNA, but the latter is believed to require complementary strand synthesis prior to successful recombination with the host genome (Kowalczykowski et al., 1994; van der Ende et al., 1983). Thus, there is no reason *a priori* why DNA must be translocated and integrated as ssDNA during NT, especially given the remarkably high rates of recombination observed for dsDNA (Babic et al., 2008). Thus, it is tempting to speculate that one reason cells integrate ssDNA during NT is to facilitate epigenetic transfer of gene products during the process. Induction of natural competence and/or successful natural transformation only occurs in a small subpopulation of cells within transformable populations, which is thought to occur as a bet-hedging strategy. This could be due to the fact that committing to the competence state incurs a fitness cost (Johnsen et al., 2009; Veening et al., 2008; Wylie et al., 2010). Alternatively, integration of tDNA from the environment, which can contain deleterious mutations, can be detrimental to the transformed cell (Moradigaravand and Engelstadter, 2013; Redfield, 1988). In our study, ~1/2 of all transformants shared tDNA-derived gene products with their untransformed siblings. The untransformed sibling that inherits these gene products obtains the potential benefit from the tDNA without having to incur the potential cost of integrating a deleterious mutation. Thus, epigenetic inheritance, as described in this study, may represent an additional layer of bet hedging to help ameliorate the fitness costs associated with horizontal gene transfer by NT.

## Acknowledgements

We gratefully acknowledge CK Ellison and NG Greene for critical comments on the manuscript. We thank CK Ellison, NG Greene, YV Brun, DK Chattoraj and ME Winkler for helpful discussions. For strains and reagents related to pneumococcal experiments, we thank DA Morrison, ME Winkler, and HT Tsui. We thank RL Gourse and the FX Barre lab for plasmids pFHC2973 and pEP74. We also thank YV Brun and members of his lab for assistance with microscopy and generous access to equipment. This work was supported by grants R35GM128674 and AI118863 from the National Institutes of Health to ABD.

## Author Contributions

ABD designed and coordinated the overall study. ABD and TND performed experiments and conducted data analysis. ABD wrote the manuscript. All authors contributed to editing the manuscript.

## Methods

### Bacterial strains and culture conditions

All *V. cholerae* strains used throughout the study were derived from the *V. cholerae* isolate E7946 (Miller et al., 1989). See Table S1 for a detailed list of all strains used in this study. All *V. cholerae* strains were engineered to be constituvely naturally competent as previously described. Strains were routinely grown at 30°C or 37°C in LB Miller agar and broth (BD Difco) supplemented with 100 µM IPTG, 200 µg/mL spectinomycin, 50 µg/mL kanamycin, 10 µg/mL trimethoprim, 100 µg/mL streptomycin, 2 µg/mL chloramphenicol, 10 µg/mL erythromycin, and/or 50 µg/mL zeocin as appropriate.

The *S. pneumoniae* strain used in this study was CP2137, Δcps ΔcomA derivative of strain Rx1 which was generously provided by Dr. Donald Morrison (Table S1). All pneumococcal strains were routinely grown in CAT+GP (CAT = 10 g/L N-Z-Amine A, 5 g/L tryptone, 1 g/L yeast extract, and 5 g/L NaCl, which was supplemented with fresh 0.2% glucose and 16mM K_2_HPO_4_) and on Trypticase Soy Agar with 5% sheep’s blood (BD BBL) supplemented with 0.3 µg/mL erythromycin and/or 250 µg/mL kanamycin as appropriate. Cultures in CAT+GP were grown statically at 37°C, while agar plates were incubated at 37°C in candle extinction jars.

### Construction of mutant strains

All *V. cholerae* and *S. pneumoniae* mutant strains were generated by SOE PCR and natural transformation, cotransformation, or MuGENT exactly as previously described (Dalia, 2018; Dalia et al., 2014; Dalia et al., 2017). For a list of all primers used in strain construction see Table S2. All mutants were confirmed by PCR and/or sequencing. The orthologous parB/S systems (yGFP-Δ23ParB_MT1_/parS_MT1_ = yGFP-ParB1/*parS1*; CFP-Δ30ParB_P1_/parS_P1_ = CFP-ParB2/*parS2*) were amplified from pFHC2973 (Nielsen et al., 2006)(*P_lac_-CFP-parB_P1_ yGFP-parB_MT1_*) and pEP74 (*P_lac_-yGFP-parB_MT1_*) and integrated into the *lacZ* locus using the primers indicated in Table S1. The yGFP gene in the yGFP-parB1 constructs from these sources was replaced with mCherry or CyPet as indicated in Table S1 using the primers indicated in Table S2.

### Imaging and analysis

Unless otherwise indicated, cell were prepared by growing cells to late log in LB supplemented with 20 mM MgCl_2_, 10 mM CaCl_2_, and 100 µM IPTG. Cells were then washed once and resuspended in instant ocean medium (7 g/L; Aquarium Systems). Cells were then mixed with tDNA as indicated and placed under 0.2% gelzan pads made with instant ocean medium supplemented with 100 µM IPTG unless otherwise indicated. Phase contrast and fluorescence images were collected on a Nikon Ti-2 microscope using a Plan Apo ×60 objective, a YFP, CFP, and/or mCherry filter cube, a Hamamatsu ORCAFlash4.0 camera and Nikon NIS Elements imaging software. Cell numbers were quantified using MicrobeJ (Ducret et al., 2016). Time-lapse imaging was performed with an interval of between 1 min and 10 min as indicated.

For tracking ComM foci, rare cells that started with a focus in timelapses (usually <<1%) were excluded from the analysis. Focus duration was determined by assessing the number of frames that ComM foci were evident. If ComM foci were separated by 2 or more frames they were considered independent events (i.e. scored as two ComM foci if there were two intervening frames that lacked foci). Based on the average duration of ComM foci being ~9 mins (Fig. 1, assessed via 1-min intervals), subsequent experiments were imaged at a 3-min intervals to allow for longer time lapses and to minimize photobleaching. To assess the percentage of cells that formed a ComM focus, the number of cells that produced at least one ComM focus within the duration of the time lapse was divided by the total number of cells analyzed. As a result, experiments that tracked cells for a shorter period of time yielded a lower percentage of cells that exhibited a ComM focus, while longer duration time lapses yielded a higher percentage of cells that exhibited a ComM focus (i.e. compare Fig. 1 = 45 min time lapse to Fig. 2 = 5 hour time lapse).

Successful DNA integration using integration of a *parS1* site, was assessed by tracking the formation of a *de novo* CyPet-ParB1 focus within a cell via time-lapse microscopy. All ComM foci formed prior to the appearance of the CyPet-ParB1 focus were scored as independent attempts at DNA integration. The success rate was defined both on a ‘per cell’ basis and on a ‘per attempt’ basis. For the per cell basis, this was defined as the percentage of the cells that succeeded to integrate the *parS1* site into their genome (which was observed as formation of a de novo CyPet-ParB1 focus in the cell) relative to the total number of cells that attempted to integrate DNA (i.e. the number of cells that generated at least one ComM focus). For the per attempt basis, this was defined as the percentage of cells that succeeded to integrate the parS1 site into their genome (which was observed as formation of a de novo CyPet-ParB1 focus in the cell) relative to the total number of attempts at DNA integration observed (i.e. the total number of ComM foci observed. Those cells that succeeded to integrate the *parS* site were demarcated “Success”, while those cells that attempted to integrate tDNA (i.e. formed a ComM focus) but ultimately failed to integrate the *parS* site were demarcated “Fail”. The number and duration of ComM foci within cells of each class were analyzed as indicated above and plotted.

To assess ssDNA vs dsDNA integration, we performed time-lapse microscopy. Only cells that initially contained one genome (i.e. started with only a single *parS2* site) and formed a de novo *parS1* site during the time lapse were analyzed. Cells that generated predivisional cells with both a ParB1 and ParB2 focus were scored as having undergone ‘ssDNA integration’, while those that formed two ParB1 foci were scored as having undergone ‘dsDNA integration’.

The time required for expression of tDNA-derived GFP following chromosome replication was determined by time-lapse microscopy. The integration of tDNA resulted in ablation of the *parS1* site and corresponding ParB1 focus. However, upon chromosome replication, the *parS1* site is restored and cells reform the corresponding ParB1 focus (as shown in Fig. 4B). Thus, the reappearance of the ParB1 focus was used to demarcate chromosome replication. The time required for tDNA expression was then defined as the first frame following chromosome replication in which GFP expression was evident. Occasionally, cells expressed GFP without reappearance of the ParB1 focus, which likely represented events where dsDNA integration had occurred. We could not demarcate *gfp* expression relative to chromosome replication reliably in these events; therefore, these cells were excluded from the analysis.

Epigenetic inheritance of antibiotic resistance was tracked via time-lapse microscopy. Cells containing a chromosomally inactivated *gfp* allele (*P_tac_*-*gfp\**::*parS1*) were transformed with DNA that would repair the *gfp* gene and contained a linked Ab^R^ marker and *parS2* site. The order of these 3 markers on the tDNA was *parS2*-Ab^R^-*P_tac_*-*gfp*. This order was critical because the first and last markers (*parS2* and *P_tac_*-*gfp*) could be visually tracked by formation of a ParB2 focus and GFP expression, respectively. So, if cells received the first and last markers, it was assumed that they would also have integrated the intervening Ab^R^ marker. Thus, only cells that received both the first and last markers were analyzed for epigenetic antibiotic inheritance. In this assay, cells were first incubated with tDNA for 3 hours in instant ocean medium and were then placed under an 0.2% gelzan pad made in M9 minimal medium supplemented with 1% glucose and the appropriate antibiotic (kanamycin 100 µg/mL, erythromycin 10 µg/mL, or chloramphenicol 2 µg/mL as indicated). Cells were imaged every 5-10 mins for up to 12 hours.

### Natural transformation / antibiotic challenge assays

For a schema of the experimental procedure see Fig. S4A-B. For experiments with *V. cholerae*, cells were grown overnight in LB supplemented with 100 µg/mL IPTG. Then, ~10^8^ cells were washed, resuspended, and diluted into instant ocean medium. Next, ~3 µg of tDNA (Kan^R^ or Spec^R^ as appropriate) was added to reactions, while nothing was added to the ‘no DNA’ control reactions. Reactions were incubated at 30°C for 5 hours to allow for DNA integration. DNAase I (NEB) was then added to reactions to eliminate all remaining extracellular DNA. Next, reactions were diluted into LB medium and outgrown for the indicated amount of time (0 min – 240 min). Following outgrowth, cells were washed twice and resuspended in instant ocean medium. Next, reactions were challenged with a high dose of antibiotic (kanamycin 1000 µg/mL or spectinomycin 10,000 µg/mL as appropriate) for 3 hours to kill all susceptible cells. Cells were then washed twice, resuspended in instant ocean medium, and plated for quantitative culture as indicated to determine the number of transformants, non-transformants, and illegitimate recombinants that survived the antibiotic treatment.

For experiments with *S. pneumoniae*, cells were grown overnight on blood agar plates. The following day, cells were suspended off of these plates with CAT+GP and diluted into 10 mL of CAT+GP and grown statically at 37°C for 3-4 hours. Then, ~1mL transformation reactions were prepared with the following: 100 µL CAT+GP, 50 µL 20 mM CaCl_2_, 2.5 µL 250 µg/mL CSP, and 850 µL of cells adjusted to an OD_600_ = 0.05. Next, ~900 ng of tDNA (Δ*nanB*::Kan^R^) was added to ‘+DNA’ reactions, while nothing was added to the ‘NO DNA’ reactions. Transformation reactions were incubated at 37°C statically for 30 min to allow for DNA integration. Then, DNase I was added to reactions to digest any uningested DNA. Reactions were incubated at 37°C statically for an additional 30 min. Reactions were then outgrown in CAT+GP medium for the indicated amount of time (0 min - 360 min). Following outgrowth, cells were washed once and resuspended in instant ocean medium containing kanamycin at a final concentration of 10,000 µg/mL. Reactions were incubated at 37°C statically for 3.5 hours to kill susceptible cells. Cells were then pelleted, washed twice, resuspended in instant ocean medium, and plated for quantitative culture as indicated to determine the number of transformants (Kan^R^ CFUs), non-transformants (Erm^R^ CFUs), and illegitimate recombinants (Kan^R^+Erm^R^ CFUs) that survived the kanamycin treatment.

## Supplementary Materials

**Fig. S1.**
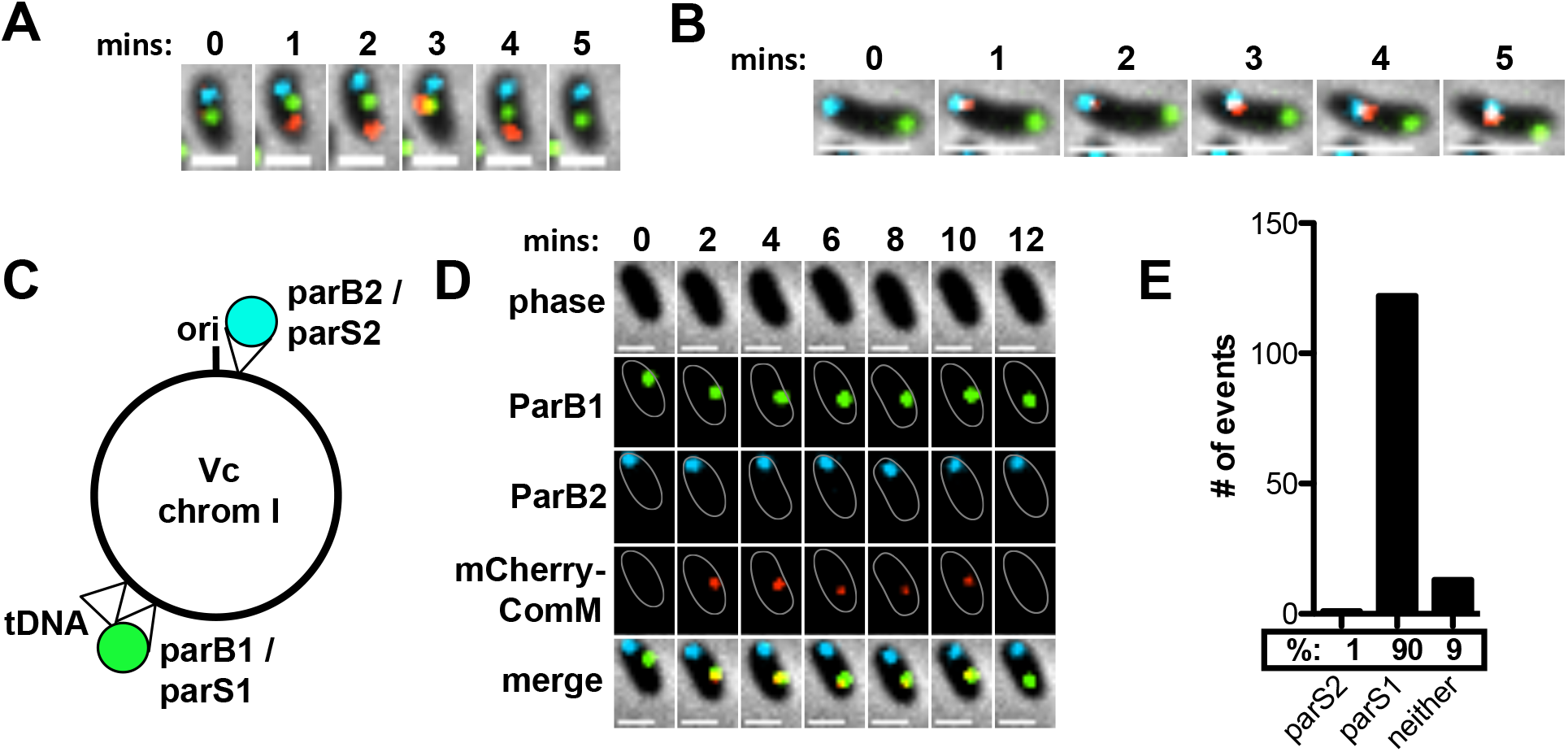
ComM fluorescent fusions serve as a marker for homologous recombination during NT, Related to Figure 1. (**A** and **B**) Montages of timelapse imaging. Cells constitutively expressed yGFP-ParB1/CFP-ParB2, contained a *parS2* site proximal to the origin, a *parS1* site proximal to the terminus, and expressed mCherry-ComM. Cells were incubated with tDNA that should integrate proximal to the *parS2* site (see Fig. 1D for a schematic). The montage in **A** shows a cell where the mCherry-ComM focus does not overlap with the *parS2* site. Scale bar, 1 µm. The montage in **B** shows a cell where the mCherry-ComM focus displays colocalized movement with the *parS2* site. Scale bar, 2 µm. (**C**) Schematic, (**D**) example montage of timelapse imaging, and (**E**) quantification of colocalization for an experiment where cells were transformed with tDNA that integrates proximal to the *parS1* site instead. Scale bar in **D** is 1 µm. Data are from two independent experiments; n = 136 ComM foci analyzed.

**Fig. S2.**
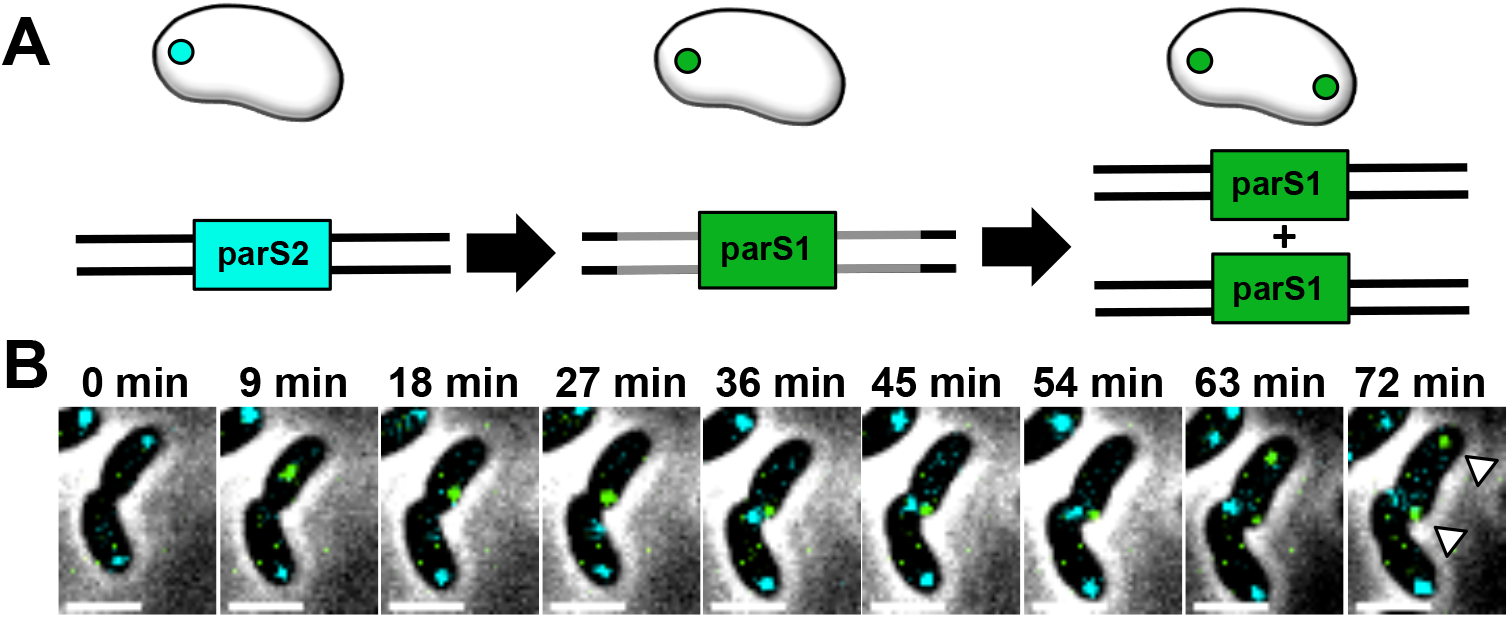
Example of dsDNA integration during NT, Related to Figure 3. Cells constitutively expressed yGFP-ParB1/CFP-ParB2 and contain a *parS2* site in the genome. Cells were then transformed with tDNA that would replace the *parS2* site with a *parS1* site. (**A**) Schematic to indicate the experimental setup and expected results for dsDNA integration. (**B**) Montage of timelapse imaging for dsDNA integration during NT. After integration, chromosome replication and segregation yields two yGFP-ParB1 foci (white arrows), which is consistent with dsDNA integration. Scale bar, 2µm.

**Fig. S3.**
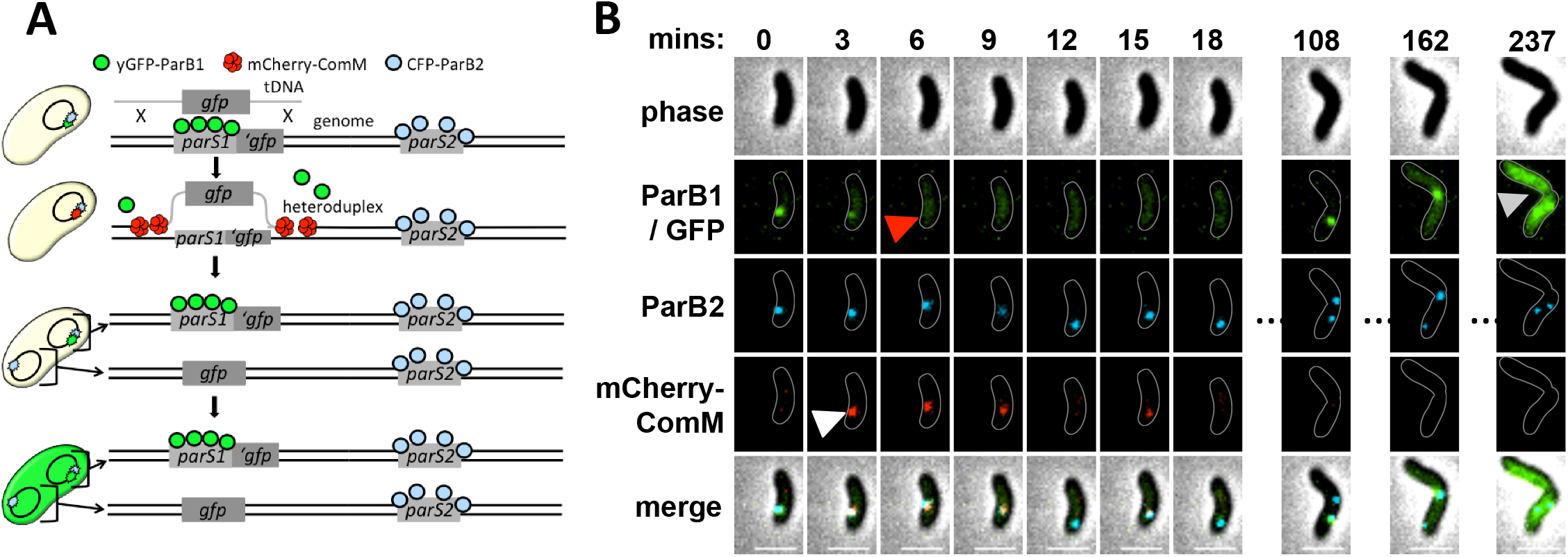
Deletion of an established *parS* site provides a sensitive and immediate readout for tDNA integration in single cells, Related to Figure 2. (**A**) Schematic indicating the experimental setup and proposed steps of tDNA integration. Cells constitutively expressed yGFP-ParB1 and CFP-ParB2, contained *parS1* and *parS2* sites in close proximity in the genome, and expressed mCherry-ComM. The *parS1* site disrupted a chromosomally integrated *gfp* gene. Cells were transformed with tDNA to delete the *parS1* site and restore the *gfp* gene. Experiments were imaged every 3 min for 5 hours. (**B**) Montage of timelapse imaging of a representative cell that successfully integrates the tDNA. Formation of a ComM focus (white arrow) immediately precedes tDNA integration, which is observed as the loss of the yGFP-ParB1 focus (red arrow). Successful DNA integration is also confirmed by elevated GFP expression (gray arrow). Scale bars, 2µm.

**Fig. S4.**
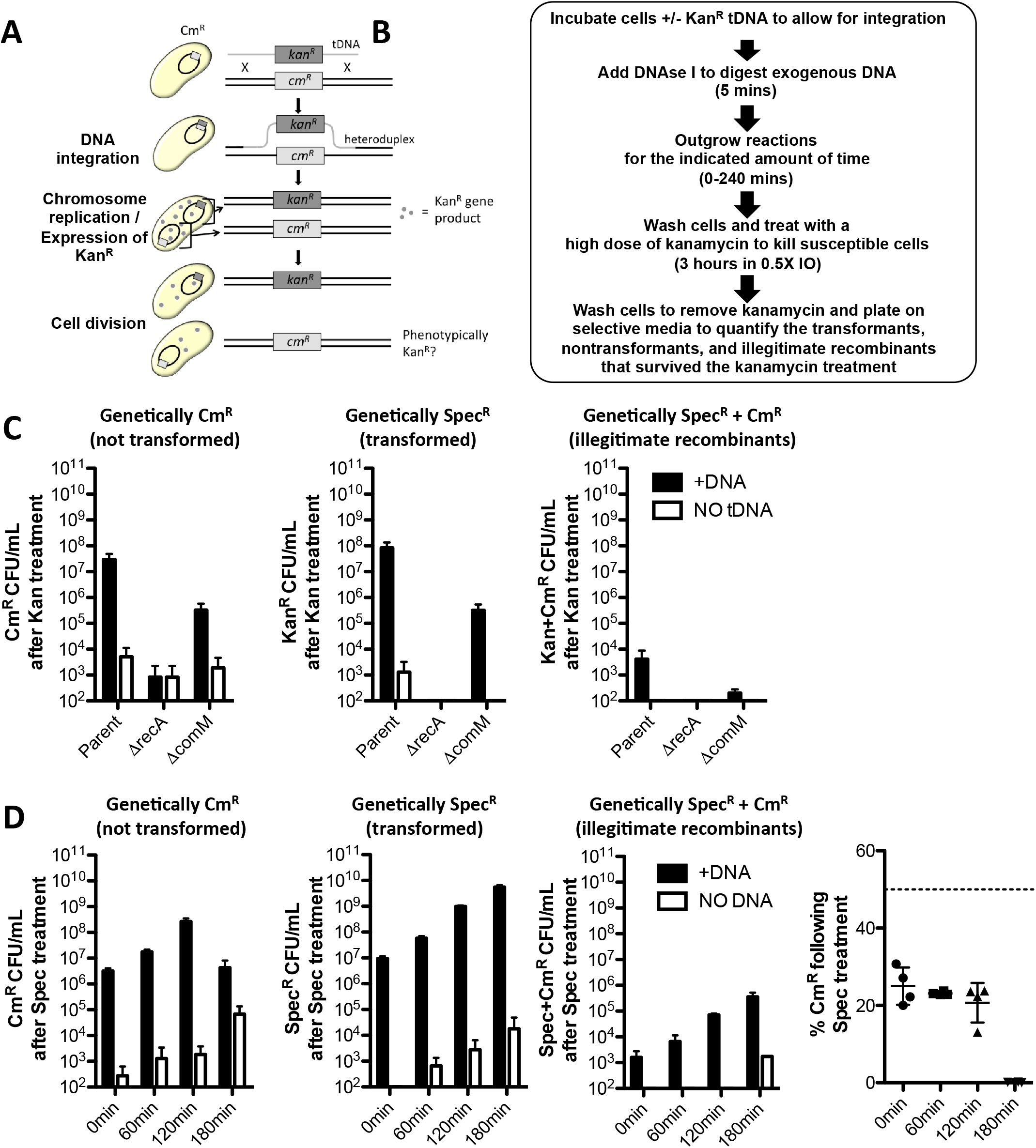
Epigenetic inheritance of antibiotic resistance requires DNA integration and is relevant to other aminoglycosides, Related to Figure 5. (**A-B**) Schema for testing transgenerational epigentic inheritance of antibiotic resistance. (**C**) Epigenetic inheritance of Kan^R^ was tested in the indicated mutant strains following 60 mins of outgrowth. (**D**) Epigenetic inheritance of antibiotic resistance was tested with Spec^R^ tDNA. Cells were outgrown for the amount of time indicated on the X-axis prior to treatment with a lethal dose of spectinomycin to kill susceptible cells. All data are shown as the mean ± SD and are from 4 independent experiments.

**Fig. S5.**
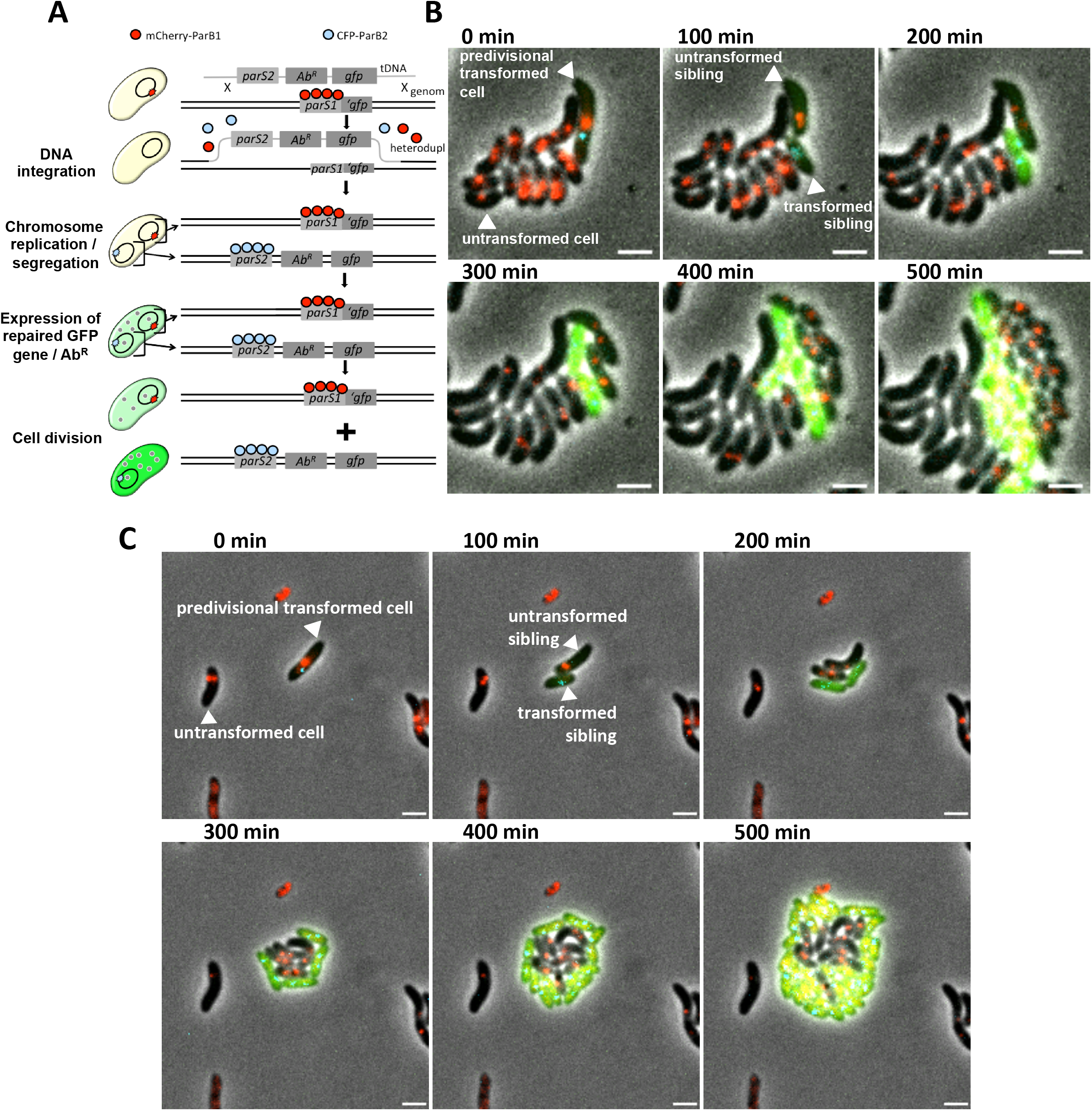
Epigenetic inheritance during NT promotes transgenerational resistance to diverse classes of antibiotics, related to Figure 6. (**A**) Schema of the experimental approach used to test epigenetic inheritance of antibiotic resistance as in Fig. 6 for kanamycin and here for (**B**) erythromycin and (**C**) chloramphenicol. Cells were transformed with Erm^R^ and Cm^R^ tDNA, respectively, and grown under pads containing the corresponding antibiotic (10 µg/mL erythromycin or 2 µg/mL chloramphenicol). Compared to ‘untransformed cells’, which do not grow, the ‘untransformed siblings’ (which are genetically Ab^S^) grow and divide for a number of generations in the presence of the antibiotic. This indicates that the untransformed sibling likely epigenetically inherited the Ab^R^ gene product from its transformed sibling. Experiments were imaged every 10 min for 12 hours. Scale bars, 2 µm.

### Supplementary Tables

**Table S1.** Strains used in this study

| Strains used in the indicated Figures |  |  |  |
| --- | --- | --- | --- |
| Use | Relevant figure(s) | Full genotype and antibiotic resistances | Reference / (strain#) |
| Parent used to generate all of the *V. cholerae* strains for this study | N/A | E7946 Sm<sup>R</sup>, $\Delta lacZ::lacI_q$, $P_{tac}-tfoX$, $\Delta luxO::Spec^R$, *comEA-mCherry*, $\Delta VC1807::Kan^R$, *pilA* S67C | (Ellison et al., 2018) |
| Tracking GFP-ComM foci | Fig. 1A-C | E7946 Sm<sup>R</sup>, $P_{tac}-tfoX$, $\Delta luxO::Spec^R$, *comEA-mCherry*, $\Delta VC1807::Cm^R$, *pilA* S67C, 0.11 Mbp::*parS*<sub>MT1</sub>, *gfp-comM*, $\Delta lacZ::P_{lac}-cyPet-parB_{MT1}$ Zeo<sup>R</sup> | This study / SAD1924 |
| Colocalizing mCherry-ComM and marked genomic loci | Fig. 1D-F, Fig. S1A-E | E7946 Sm<sup>R</sup>, $P_{tac}-tfoX$, $\Delta luxO$, $\Delta VC1807::Cm^R$, *pilA* S67C, *mCherry-comM*, 0.11 Mbp::*parS*<sub>P1</sub>, 1.963 Mbp::*parS*<sub>MT1</sub>, $\Delta lacZ::P_{lac}-CFP-parB_{P1}$ yGFP-*parB*<sub>MT1</sub> Zeo<sup>R</sup> | This study / TND1379 (SAD2440) |
| Quantifying success rates for integration of $\Delta 0kb::parS1$ | Fig. 2A-F | E7946 Sm<sup>R</sup>, $P_{tac}-tfoX$, $\Delta luxO::Spec^R$, *comEA-mCherry*, $\Delta VC1807::Kan^R$, *pilA* S67C, *gfp-comM*, $\Delta lacZ::P_{lac}-cyPet-parB_{MT1}$ Zeo<sup>R</sup> | This study / SAD1923 |
| Quantifying success rates for integration of $\Delta 1.5kb::parS1$ | Fig. 2B-H | E7946 Sm<sup>R</sup>, $P_{tac}-tfoX$, $\Delta luxO::Spec^R$, *comEA-mCherry*, $\Delta VC1807::Kan^R$, *pilA* S67C, *gfp-comM*, $\Delta lacZ::P_{lac}-cyPet-parB_{MT1}$ Zeo<sup>R</sup>, 0.11 Mbp::*Cm*<sup>R</sup> | This study / TND1435 (SAD2441) |
| Quantifying ssDNA vs dsDNA integration | Fig. 3A-C, Fig. S2, movie S1 | E7946 Sm<sup>R</sup>, $P_{tac}-tfoX$, $\Delta luxO::Spec^R$, *comEA-mCherry*, $\Delta VC1807::Cm^R$, 0.11 Mbp::*parS*<sub>P1</sub>, *pilA* S67C, $\Delta lacZ::P_{lac}-CFP-parB_{P1}$ yGFP-*parB*<sub>MT1</sub> Zeo<sup>R</sup> | This study / SAD1950 |
| Colocalizing DNA integration and ComM foci | Fig. S3A-B | E7946 Sm<sup>R</sup>, $P_{tac}-tfoX$, $\Delta luxO$, *mCherry-comM*, 1.963 Mbp::*parS*<sub>P1</sub>, $\Delta VC1807::Cm^R-P_{tac}-gfp^*::parS_{MT1}$, *pilA* S67C, $\Delta lacZ::P_{lac}-CFP-parB_{P1}$ yGFP-*parB*<sub>MT1</sub> Zeo<sup>R</sup> | This study / TND1307 (SAD2438) |
| Tracking epigenetic inheritance of GFP | Fig. 4A-C, movie S2 | E7946 Sm<sup>R</sup>, $P_{tac}-tfoX$, $\Delta luxO$, *pilA* S67C, 1.963 Mbp::*parS*<sub>P1</sub>, $\Delta VC1807::Cm^R-P_{tac}-gfp^*::parS_{MT1}$, $\Delta lacZ::P_{lac}-CFP-parB_{P1}$ *mCherry-parB*<sub>MT1</sub> Zeo<sup>R</sup> | This study / TND1338 (SAD2439) |
| Testing epigenetic inheritance of antibiotic resistance via natural transformation / antibiotic challenge | Fig. 5A-D, Fig. S4A-D | E7946 Sm<sup>R</sup>, $P_{tac}-tfoX$, $\Delta luxO$, *pilA* S67C, *comEA-mCherry*, $\Delta lacZ::lacI_q$, $\Delta VC1807::Cm^R$ | This study / TND0904 (SAD2436) |
| Testing epigenetic inheritance of kanamycin / erythromycin resistance via time lapse microscopy | Fig. 6, movies S3-S4, Fig. S5A-B | E7946 Sm<sup>R</sup>, $P_{tac}-tfoX$, $\Delta luxO$, *pilA* S67C, $\Delta VC1807::Cm^R-P_{tac}-gfp^*::parS_{MT1}$, $\Delta lacZ::P_{lac}-CFP-parB_{P1}$ *mCherry-parB*<sub>MT1</sub> Zeo<sup>R</sup> | This study / TND1449 (SAD2442) |
| Testing epigenetic inheritance of chloramphenicol resistance via time lapse microscopy | Fig. S5A-C, movie S5 | E7946 Sm<sup>R</sup>, P<sub>tac</sub> - *tfoX*, $\Delta luxO$ , *pilA* S67C, $\Delta VC1807::Erm^R$ -P<sub>tac</sub> - *gfp* *:: *parS<sub>MT1</sub>*, $\Delta lacZ::P_{lac}$ -CFP- *parB<sub>P1</sub>* mCherry- *parB<sub>MT1</sub>* Zeo<sup>R</sup> | This study / TND1458 (SAD2443) |
| Parent strain used to generate all of the *S. pneumoniae* strains for this study. | N/A | Rx1 *hex malM511 str-1 bgl-1*; low-α-galactosidase background; Sm<sup>R</sup>, $\Delta cps$, $\Delta cheshire$ *comA*<sup>-</sup> Cm<sup>R</sup> | (Weng et al., 2009) |
| *S. pneumoniae* host for testing epigenetic inheritance of antibiotic resistance via natural transformation / antibiotic challenge | Fig. 5E-H | Rx1 *hex malM511 str-1 bgl-1*; low-α-galactosidase background; Sm<sup>R</sup>, $\Delta cps$, $\Delta cheshire$ *comA*<sup>-</sup> Cm<sup>R</sup>, $\Delta nanB::Erm^R$ | This study / SAD2403 |
| **Strain gDNA used as template to amplify tDNA** |  |  |  |
| **Use(s)** | **Relevant figure(s)** | **Relevant tDNA genotype and primers used to amplify up the PCR product (see Table S2 for primer sequences)** | **Reference / (strain#)** |
| -success rates for tDNA integration<br>-testing ssDNA vs dsDNA integration | Fig. 2A-H, Fig. 3A-C, Fig. S2, movie S1 | tDNA = 0.11 Mbp::*parS<sub>MT1</sub>*<br><u>Amplified with:</u> BBC1852 / BBC1855 | This study / SAD1924 |
| -assess ComM focus duration<br>-colocalization of ComM foci with genomic loci | Fig. 1A-F, Fig. S1A-B | tDNA = 0.102 Mbp WT<br><u>Amplified with:</u> BBC1981 / BBC1984 | This study / SAD033 |
| -colocalization with genomic loci<br>-natural transformation / kanamycin challenge | Fig. S1C-E, Fig. 5A-D, Fig. S4C | tDNA = $\Delta VC1807::Kan^R$<br><u>Amplified with:</u> BBC1881 / BBC1882 | This study / SAD034 |
| -natural transformation / spectinomycin challenge | Fig. S4D | tDNA = $\Delta VC1807::Spec^R$<br><u>Amplified with:</u> BBC1881 / BBC1882 | This study / SAD033 |
| -natural transformation / kanamycin challenge in *S. pneumoniae* | Fig. 5E-H | tDNA = $\Delta nanB::Kan^R$<br><u>Amplified with:</u> BBC2853 / BBC2856 | This study / BBC2402 |
| -colocalize DNA integration and ComM foci<br>-tracking epigenetic inheritance of GFP | Fig. S3A-B, Fig. 4A-C, movie S2 | tDNA = $\Delta VC1807::Cm^R$-P<sub>tac</sub>-*gfp*<br><u>Amplified with:</u> BBC1881 / BBC1882 | This study / TND1166 (SAD2437) |
| -Time lapse microscopy to test epigenetic inheritance of kanamycin resistance | Fig. 6, movie S3 | tDNA = $\Delta VC1807::parS_{P1}$-*Kan*<sup>R</sup>-P<sub>tac</sub>-*gfp*<br><u>Amplified with:</u> BBC1881 / BBC1882 | This study / TND1513 (SAD2446) |
| -Time lapse microscopy to test epigenetic inheritance of erythromycin resistance | Fig. S5B, movie S4 | <u>tDNA</u> = $\Delta VC1807::parS_{P1}-Erm^R-P_{tac}-gfp$<br><u>Amplified with:</u> BBC1881 / BBC1882 | This study / TND1492 (SAD2444) |
| -Time lapse microscopy to test epigenetic inheritance of erythromycin resistance | Fig. S5C, movie S5 | <u>tDNA</u> = $\Delta VC1807::parS_{P1}-Cm^R-P_{tac}-gfp$<br><u>Amplified with:</u> BBC1881 / BBC1882 | This study / TND1498 (SAD2445) |

**Table S2.**
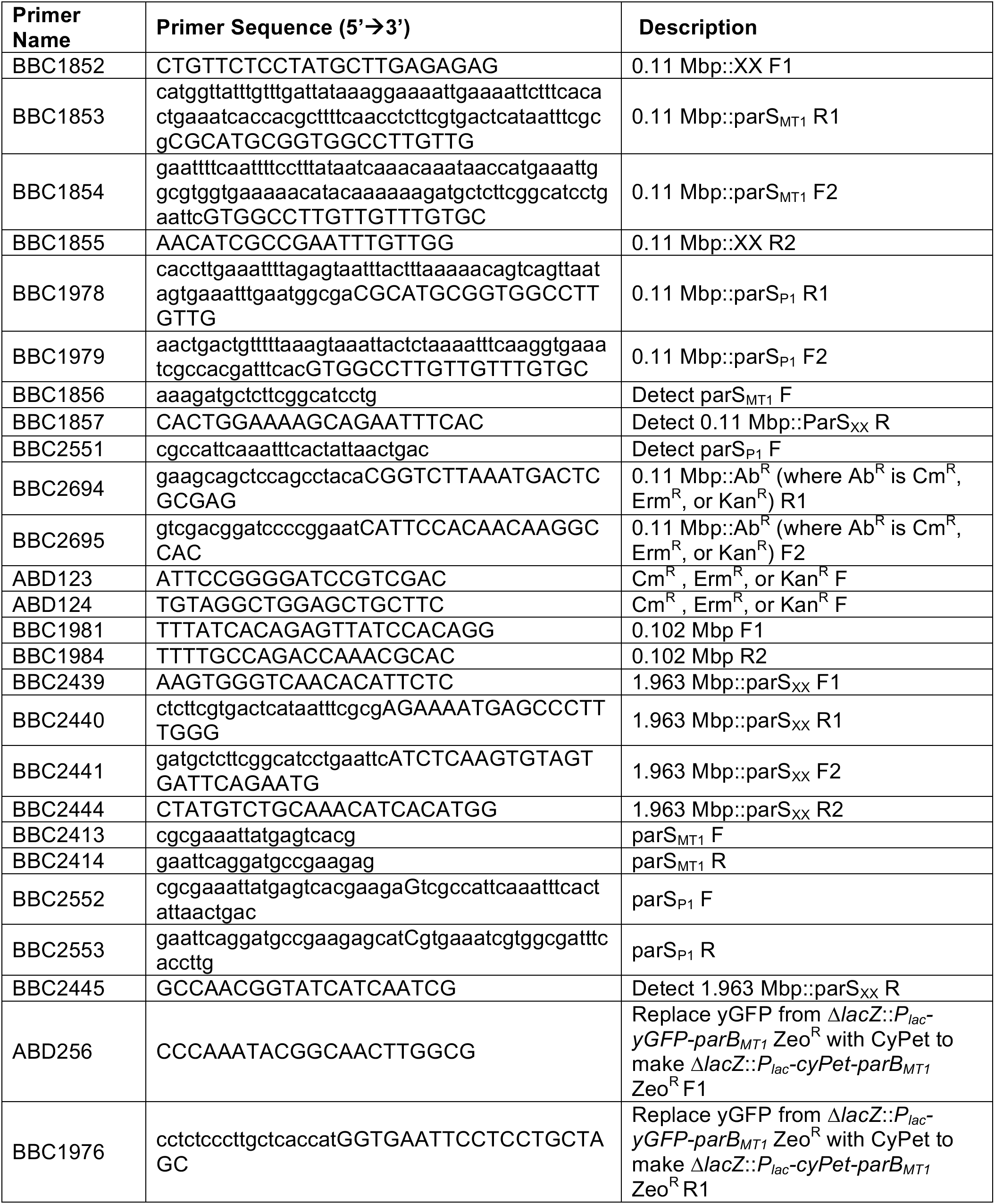
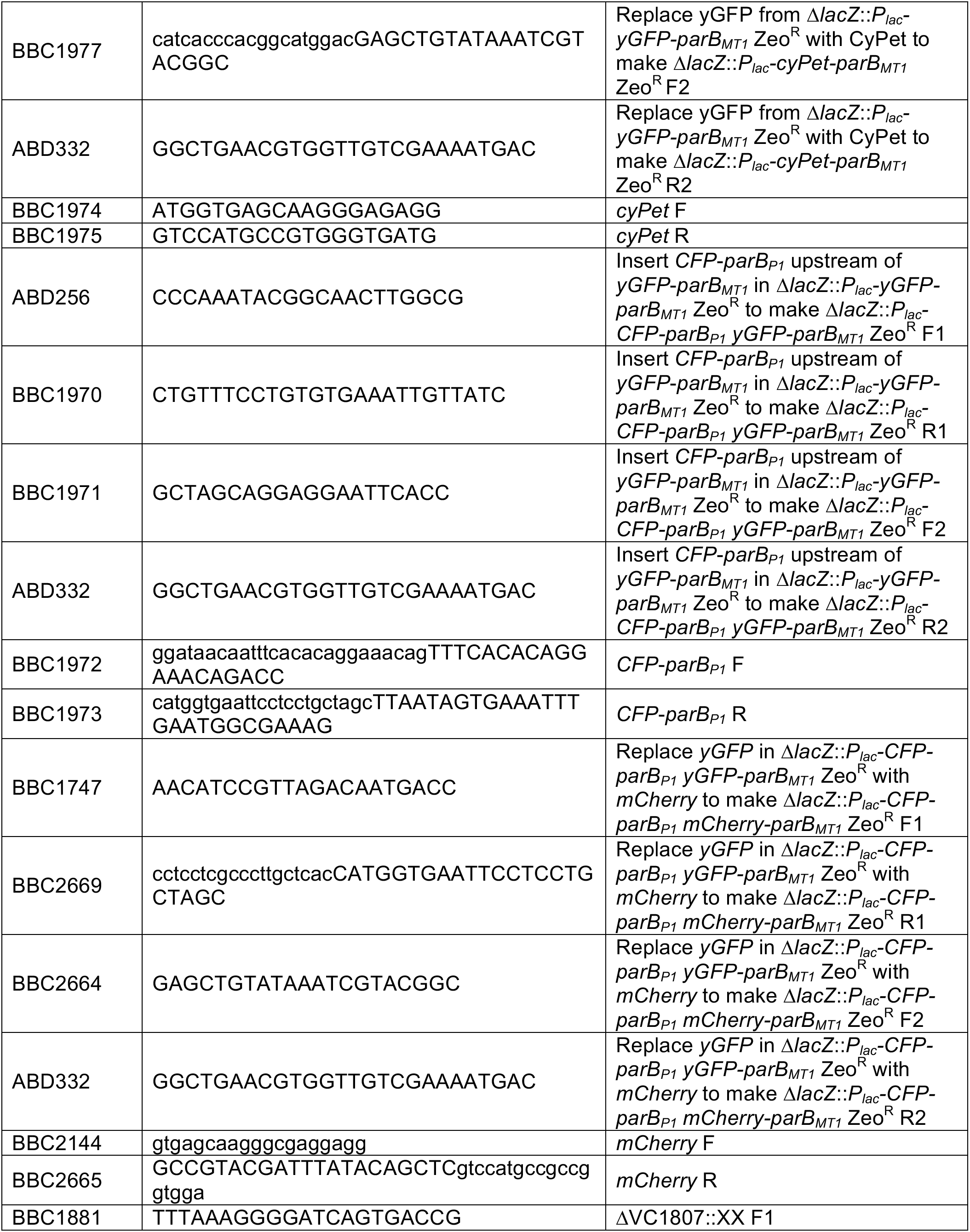
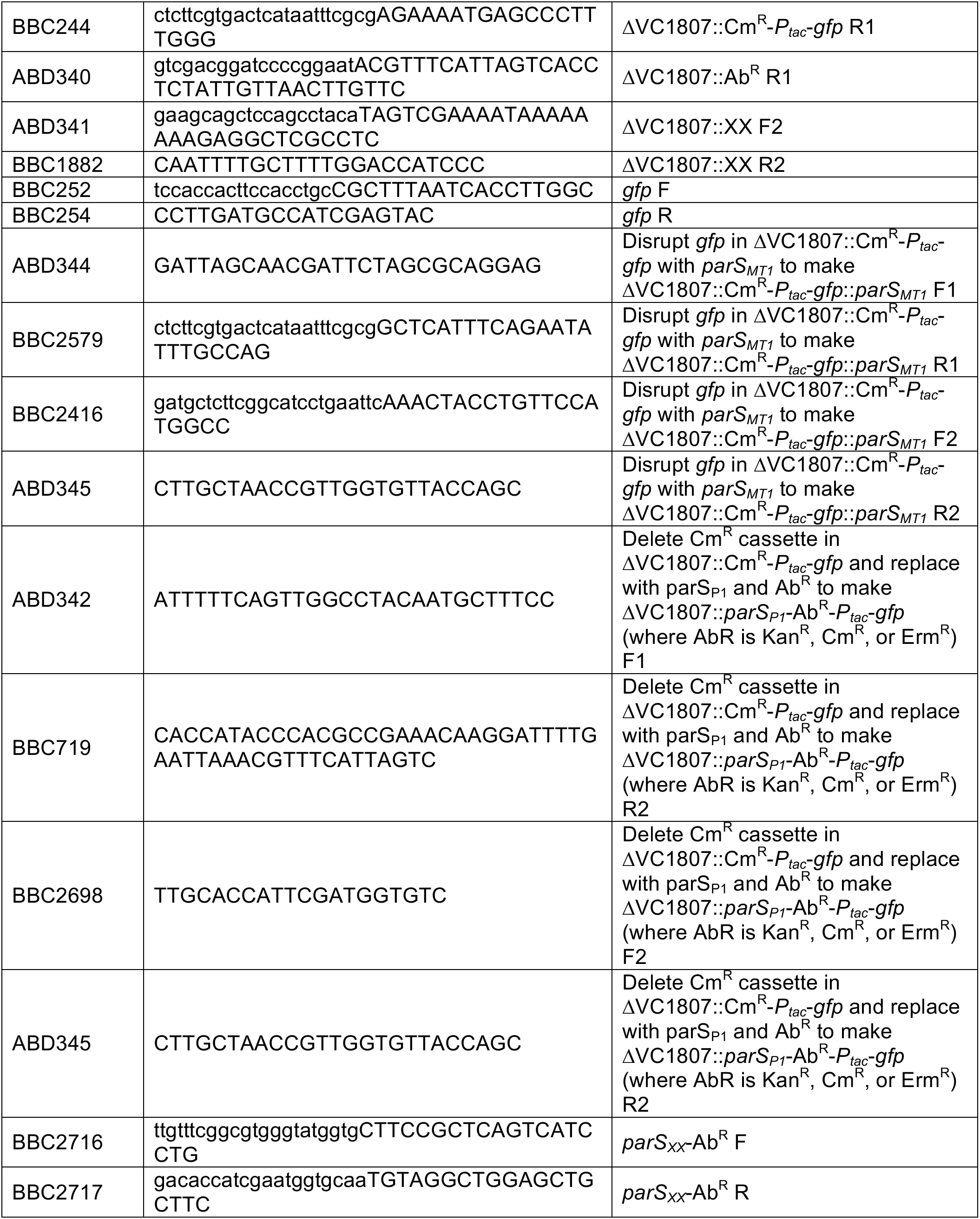
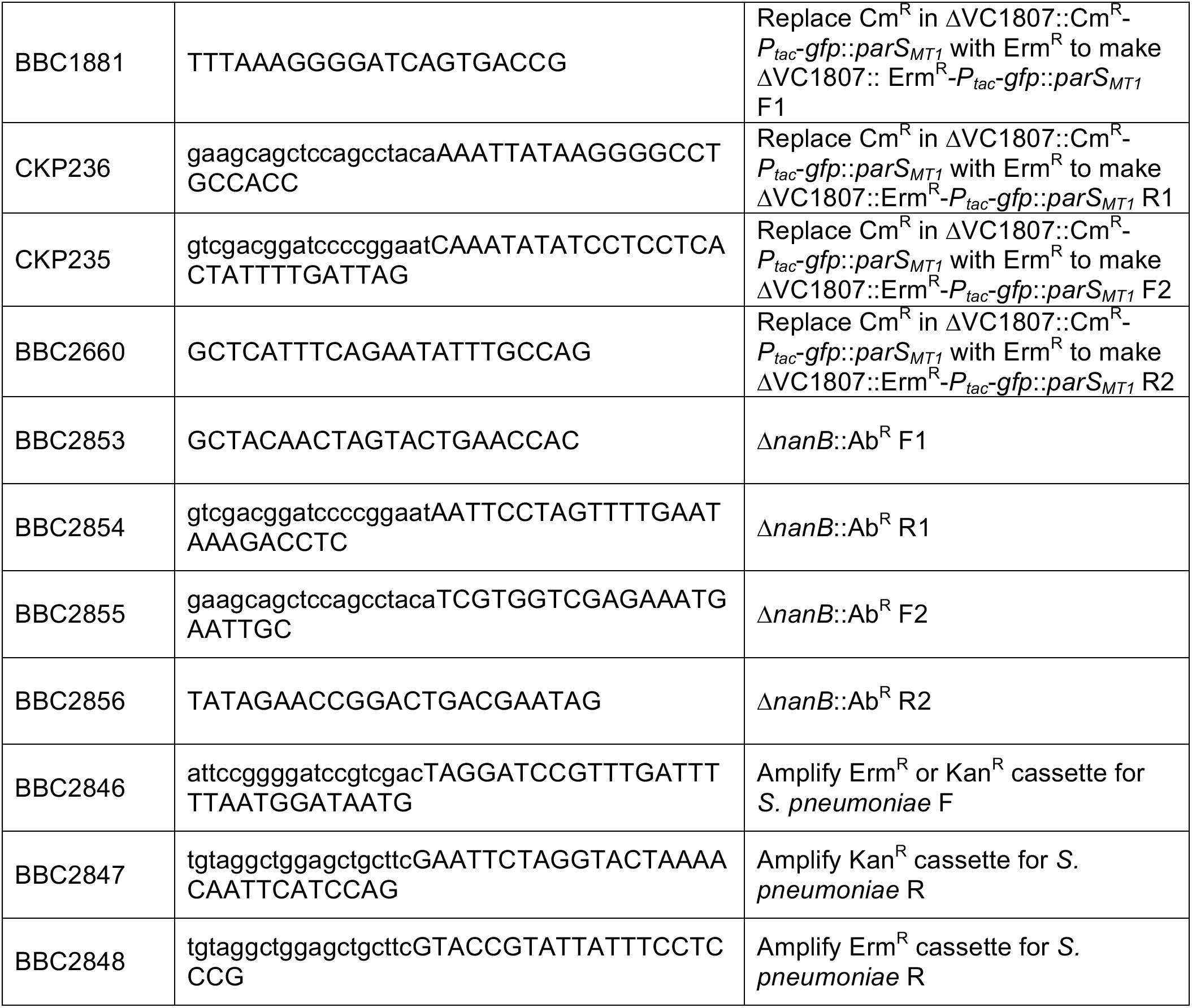
Primers used in this study

### Supplemental Movie Legends

**Movie S1 – tDNA integrates as ssDNA during NT and the resulting heteroduplex is resolved by chromosome replication.** Time lapse of the cell shown in Fig. 3B. Cells in this experiment constitutively expressed yGFP-parB1/CFP-parB2 and contained a parS2 site in the genome. Cells were then transformed with tDNA that would replace the parS2 site with a parS1 site. For a schema of the experimental setup see Fig. 3A. This experiment was repeated independently two times with similar results. The capture interval is 3 min between frames. Scale bar is 2 µm.

**Movie S2 – Epigenetic inheritance of tDNA-derived GFP during NT.** Time lapse of the cell shown in Fig. 4B. Cells in this experiment contained *parS1* (red ParB1) and *parS2* (blue ParB2) sites in close proximity on the chromosome. The *parS1* site interrupted a chromosomally integrated *gfp* gene. Cells were transformed with tDNA to delete the *parS1* site and restore the *gfp* gene. For a schema of the experimental setup see Fig. S4A. This experiment was repeated independently three times with similar results. The capture interval is 3 min between frames. Scale bar is 2 µm.

**Movie S3 – Single cell assay to demonstrate transgenerational epigenetic inheritance of kanamycin resistance during NT.** Time lapse of the cells shown in Fig. 6. Cell in this experiment constitutively expressed mCherry-ParB1/CFP-ParB2 and contained a *parS1* site in the genome that disrupts a *gfp* gene. Cells were then incubated with tDNA that would remove the *parS1* site, restore the *gfp* gene, and integrate a *parS2* site linked to a Kan^R^ marker. Imaging was performed under an M9+glucose pad containing 100 µg/mL kanamycin. For a schema of the experimental setup see Fig. S5A. This experiment was repeated independently two times with similar results. The capture interval is 5 min between frames. Scale bar is 2 µm.

**Movie S4 – Single cell assay to demonstrate transgenerational epigenetic inheritance of eythromycin resistance during NT.** Time lapse of the cells shown in Fig. S5B. Cell in this experiment constitutively expressed mCherry-ParB1/CFP-ParB2 and contained a *parS1* site in the genome that disrupts a *gfp* gene. Cells were then incubated with tDNA that would remove the *parS1* site, restore the *gfp* gene, and integrate a *parS2* site linked to an Erm^R^ marker. Imaging was performed under an M9+glucose pad containing 10 µg/mL erythromycin. For a schema of the experimental setup see Fig. S5A. This experiment was repeated independently two times with similar results. The capture interval is 10 min between frames. Scale bar is 2 µm.

**Movie S5 – Single cell assay to demonstrate transgenerational epigenetic inheritance of chloramphenicol resistance during NT.** Time lapse of the cells shown in Fig. S5C. Cell in this experiment constitutively expressed mCherry-ParB1/CFP-ParB2 and contained a *parS1* site in the genome that disrupts a *gfp* gene. Cells were then incubated with tDNA that would remove the *parS1* site, restore the *gfp* gene, and integrate a *parS2* site linked to an Cm^R^ marker. Imaging was performed under an M9+glucose pad containing 2 µg/mL chloramphenicol. For a schema of the experimental setup see Fig. S5A. This experiment was repeated independently two times with similar results. The capture interval is 10 min between frames. Scale bar is 2 µm.

